# Cellular Function of a Biomolecular Condensate Is Determined by Its Ultrastructure

**DOI:** 10.1101/2024.12.27.630454

**Authors:** Daniel Scholl, Tumara Boyd, Andrew P. Latham, Alexandra Salazar, Asma Khan, Steven Boeynaems, Alex S. Holehouse, Gabriel C. Lander, Andrej Sali, Donghyun Park, Ashok A. Deniz, Keren Lasker

## Abstract

Biomolecular condensates play key roles in the spatiotemporal regulation of cellular processes. Yet, the relationship between atomic features and condensate function remains poorly understood. We studied this relationship using the polar organizing protein Z (PopZ) as a model system, revealing how its material properties and cellular function depend on its ultrastructure. We revealed PopZ’s hierarchical assembly into a filamentous condensate by integrating cryo-electron tomography, biochemistry, single-molecule techniques, and molecular dynamics simulations. The helical domain drives filamentation and condensation, while the disordered domain inhibits them. Phase-dependent conformational changes prevent interfilament contacts in the dilute phase and expose client binding sites in the dense phase. These findings establish a multiscale framework that links molecular interactions and condensate ultrastructure to macroscopic material properties that drive cellular function.

## Introduction

Biomolecular condensates are non-stoichiometric assemblies of macromolecules that concentrate specific components while excluding others^1–3^. Their formation is driven by multivalent interactions, which can be implemented through intrinsically disordered regions (IDRs), repeated oligomerization motifs, or nucleic acid chains^4,5^. These dynamic assemblies are pervasive across biology, serving as a fundamental mechanism of cellular organization and mediating a wide array of cellular functions, including transcription regulation^6^, signal transduction^7^, and protein quality control^8,9^. They orchestrate cellular processes through the function of individual molecules within the condensate, but also through the properties of the condensate itself^10–12^. These emergent properties–arising from the collective behavior of many molecules–include viscoelasticity^13^, surface tension^14^, and electrochemical potential^15^. Despite their importance, it remains a major challenge to link condensate-level properties to the molecular-level features from which they originate.

A particularly understudied question is how condensate function relates to its internal structure^4,16^. By internal structure, we refer to stable structures on the molecular and supramolecular level that are larger than single molecules but smaller than the condensate itself. A notable supramolecular structure that has been observed for several condensates across different cellular pathways is the filament^17^. Unlike cytoskeletal filaments, which are long and designed for mechanical stability^18^, these condensate-associated filaments are typically short and flexible. The prevalence of filamentous condensates is illustrated by systems such as p62^19^, DAXX/SPOP^20^, and F-actin^21^, underscoring how useful a detailed characterization of this class of condensates could be.

To determine whether and how filamentation impacts condensate properties and function, it is essential to examine the individual and collective behavior of macromolecules across all relevant length scales–from molecular, to the filament, to the condensate level–both in dispersed and condensed states. In this study, we employed a combination of *in vivo*, *in vitro*, and *in silico* techniques, including cryo-electron tomography, single-molecule Förster resonance energy transfer (FRET), and computational simulations, to elucidate the structure-function relationship of the bacterial polar organizing protein Z (PopZ), from monomers to condensates.

PopZ^22–24^, a conserved protein within α-proteobacteria^25^, plays a crucial role in cytosol organization. First identified in *Caulobacter crescentus* (*C. crescentus)*^26–28^—a well-studied model system for asymmetric cell division—PopZ forms condensates at the ends (poles) of the cell. These condensates selectively recruit cell cycle-regulated proteins, thereby establishing signaling gradients essential for driving asymmetric cell division^29^. Deletion of the *popZ* gene in *Caulobacter* results in abnormal cell division, disrupted chromosome segregation, and the loss of cellular asymmetry^23,24^. Furthermore, aberrations in the condensate’s material properties, either too fluid or too rigid, lead to severe growth defects^30^. Beyond its role in *Caulobacter*, PopZ forms condensates across various systems, including *in vitro*, *Escherichia coli,* and human cells, and is amenable to genetic and biochemical engineering^30^.

In this study, we employed an integrative approach to decode the structural and chemical determinants of PopZ condensate formation. We uncovered a hierarchical pathway in which PopZ assembles into trimers, hexamers, filaments, and condensates. We characterized the conformational dynamics of individual PopZ molecules in both the dilute and dense phases and observed phase-dependent conformational changes. Notably, the domain responsible for client binding in the dense phase also acts as an inhibitor of condensation in the dilute phase. Finally, we perturbed specific steps in the assembly pathway to study the relationship between PopZ filaments and the properties of the resulting condensates. Our studies indicate that filamentation increases the viscosity and surface tension of PopZ condensates *in vitro* and is tightly linked to PopZ function *in vivo*. Moreover, we uncovered a simple yet sophisticated bacterial implementation of filamentous liquid condensates. While other filamentous condensates rely on separate proteins for filament formation, capping, and cross-linking, PopZ uniquely integrates all three functions into a single 77-residue helical domain at its C-terminus. This domain is regulated by an IDR that inhibits condensation in the dilute phase, ensuring proper spatial and temporal control of assembly. Our work establishes a mechanistic analysis framework to understand how molecular-level features propagate across length-scales and establishes a foundation for characterizing key attributes of filamentous condensates.

## Results

### PopZ condensates are made up of short filaments

To investigate the supramolecular structure of PopZ within condensates, we purified recombinant wild type (WT) PopZ, triggered condensation by adding 50 mM MgCl_2_, and visualized the resulting condensates by cryo-electron tomography (cryo-ET) (Fig. 1A). We observed that PopZ condensates are composed of filaments (Fig. 1A, middle). To quantitatively characterize these filaments, we segmented tomograms using the machine learning-based segmentation software DragonFly^31^ and traced individual filaments using the image analysis software Amira (Thermo Fisher) (Video S1). The distribution of filament lengths is depicted in Fig. 1B, with median and mean lengths of 34 nm and 38 nm, respectively, in agreement with a study on PopZ filaments in *Magnetospirillum gryphiswaldense*^32^. Filament length was exponentially distributed (R^2^ = 0.99), indicating that PopZ polymerization is isodesmic, in turn suggesting that the formation of filaments is not cooperative, *i.e.,* that the association constant of individual units to the filament ends is independent of filament length. In isodesmic polymers, there are no distinct nucleation and elongation phases, as the bonds between building blocks are identical and the affinity between them does not depend on their position within the filament^33,34^. Therefore, characterizing the PopZ filament reduces to identifying its building blocks and the bonds connecting them.

**Figure 1.**
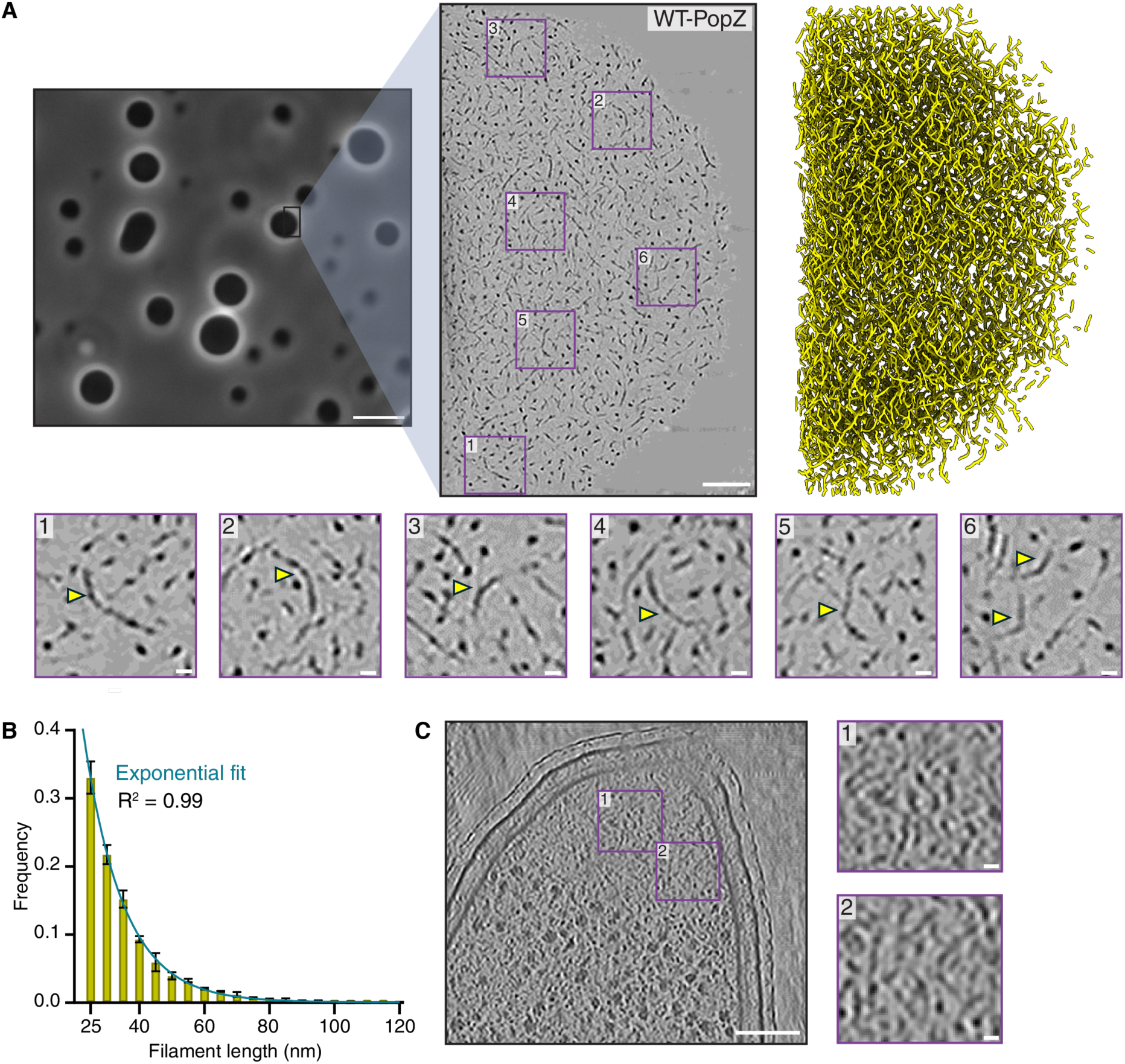
Visualization of PopZ condensates and filaments by cryo-electron tomography. **(A)** Phase-contrast imaging and cryo-ET of in vitro PopZ condensates. (top left) Phase contrast image of PopZ condensates formed by adding 50 mM MgCl2 to 5 µM of purified PopZ in 10 mM sodium phosphate (pH 7.4) with 150 mM NaCl. Scale bar: 5 µm. The zoom-in box is provided for illustrative purposes. (top middle) Cryo-ET tomogram of 2 µM purified WT-PopZ in the same buffer, captured with 53,000-fold magnification. Scale bar: 100 nm. The purple boxes indicate magnified regions of the condensate. (top right) 3D rendering of segmented tomograms. (bottom) Six zoomed-in regions illustrating the flexibility of filaments in PopZ condensates. Yellow arrows highlight filamentous structures. Scale bar: 10 nm. **(B)** Filament length distribution fits an isodesmic model. Length distribution of over 8000 filaments from three independent condensates (see Fig. S4B). Columns represent the mean filament length across the three condensates; error bars denote standard deviation. The turquoise line indicates a fit to an isodesmic model. **(C)** Cryo-ET reveals filaments within PopZ condensates in Caulobacter crescentus. Representative cryo-ET images of the PopZ microdomain in ΔpopZ Caulobacter cells expressing mCherry-tagged WT-PopZ. Scale bar: 100 nm. Filaments can be seen in the zoomed-in panel. Scale bar: 10nm. Additional reconstructions and representative filament images are shown in Fig. S1A.

To confirm that WT-PopZ forms filaments *in vivo*, we vitrified *C. crescentus* cells expressing mCherry-tagged WT-PopZ and used cryo-focused ion beam (cryo-FIB) milling to create lamellae thinner than 200 nm. This reduction in specimen thickness, combined with high-magnification cryo-ET, enabled us to achieve detailed visualization of *C. crescentus* cells. We observed the previously described PopZ microdomains at the cell poles, which were devoid of large macromolecules such as ribosomes (Figs. 1C and S1A). Inside these microdomains, we identified filamentous structures consistent with WT-PopZ filaments observed *in vitro*, although other proteins may contribute to these structures as well. Taken together, our structural characterization reveals that the underlying substructure of the PopZ condensate *in vivo* and *in vitro* consists of a meshwork of interwoven filaments, which themselves have a broad length distribution.

### The helical oligomerization domain is necessary and sufficient for filamentation and condensation

To characterize the mechanism and driving forces of PopZ condensation, we screened different salt and *p*H conditions for their ability to trigger droplet formation. Purified PopZ condenses at low micromolar concentrations upon addition of divalent cations like MgCl_2_; physiological concentrations of 2-5 mM are sufficient, while 50 mM accelerates assembly and maximizes scattering^30^. Here, we report that low *p*H (< 6.0) or increased concentrations of salt (> 300 mM NaCl) also promote condensation (Fig. S1B). We tested three buffering agents−sodium phosphate, HEPES, and Tris-HCl−all of which supported droplet formation. PopZ formed smaller droplets in Tris-HCl buffer and occasionally formed non-spherical, aggregate-like clusters in a buffer lacking both salt and divalent cations (Fig. S1B, upper rightmost panel). Using a physiologically relevant buffer, with lower magnesium concentration and a crowding agent (10 mM sodium phosphate *p*H 7.4 with 150 mM NaCl, 1 mM MgCl_2_, and 5% PEG), we observed robust droplet formation (Fig. S1B, bottom rightmost panel).

We speculated that conditions facilitating condensation highlight the significance of PopZ’s intrinsically disordered region (IDR). PopZ has a modular structure consisting of a C-terminal helical oligomerization domain (OD) necessary for condensation^29,35,36^; a negatively charged, proline-rich IDR that imparts fluidity to the condensates^30^; and an N-terminal helical region (H1) that recruits binding partners into the microdomain^37,38^ (Fig. 2 A-B). Because divalent cations, increased salt concentrations, and lower *p*H all promoted droplet formation, we hypothesized that these conditions screen repulsive interactions driven by the negatively charged IDR. We have previously shown that the length and charge distribution of the IDR affect the material properties of PopZ condensates *in vivo*^30^, indicating that the IDR influences condensate function.

**Figure 2.**
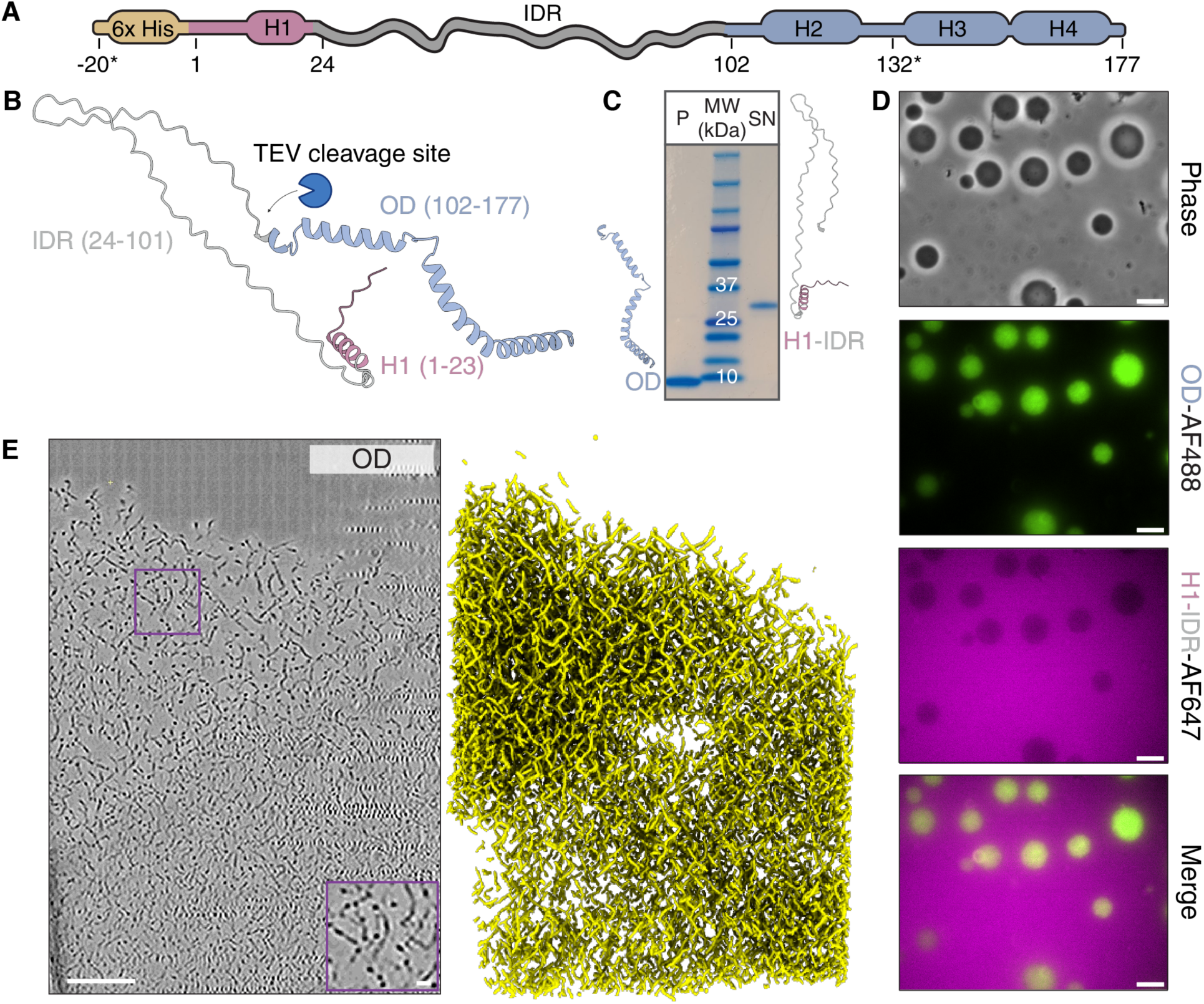
The oligomerization domain is necessary and sufficient for filamentation and condensation. **(A)** Schematic of the PopZ construct. The construct includes an N-terminal 6xHIS fusion (brown) fused to PopZ from *Caulobacter crescentus*, which codes for a short helical N-terminal region, H1 (pink box), a 78 amino-acid intrinsically disordered region (IDR, gray curly line), and a C-term region comprising three predicted helices: H2, H3, H4 (blue boxes). Positions -20 and 132 (indicated by asterisks) were used to covalently attach fluorescent dyes for imaging and smFRET studies. **(B)** AlphaFold 2 model of the PopZ monomer shown for illustrative purposes. A TEV cleavage site, introduced between residues 99 and 100, is present in our PopZ construct only when specified. **(C-D)** TEV cleavage separates the H1-IDR region from the OD, leading the OD to condense and H1-IDR to be excluded from the condensates. **(C)** SDS-PAGE gel revealed with Coomassie staining. TEV protease was added to TEV-cleavable PopZ *in vitro* and incubated for one hour before pelleting down the dense phase (2000 g, 3 min). P is the pellet fraction, which contains the OD, SN is the supernatant, which contains the IDR. The IDR migrates as a larger species due to its stretches of densely packed negative charges. **(D)** Phase and fluorescence images 30 minutes after incubation with TEV protease. The formed droplets (phase) are composed of cleaved OD (OD-AF488) and exclude cleaved H1-IDR (H1-IDR-AF647). Scale bars: 5 µm **(E)** OD condensates are made up of filaments. (left) cryo-ET tomogram of 5 µM H1-IDR-TEV-OD cleaved for 3 minutes with TEV protease in 20 mM Tris-HCl *p*H 8.0, 0.5 mM EDTA, 2 mM DTT. Magnification: 53,000x. The purple box indicates a zoomed-in region of the condensate. Scale bar: 100 nm. (right) 3D rendering of filaments of the segmented tomograms.

To test the IDR’s role in condensation, we designed a construct with a TEV protease cleavage site between the IDR and the OD (H1-IDR-TEV-OD). After purification, this construct did not form droplets until we added MgCl_2_, mirroring the behavior of WT-PopZ. Likewise, H1-IDR-TEV-OD did not form droplets in a TEV cleavage buffer (20 mM Tris HCl *p*H 8.0, 0.5 mM EDTA, 2 mM DTT). Upon addition of the TEV protease and initiation of cleavage, which separated H1-IDR from the OD, droplets formed within seconds. We then pelleted these droplets by centrifugation and analyzed both the pellet and the supernatant using SDS-PAGE. The gel revealed that the droplets were enriched in the OD, whereas the IDR remained in the supernatant (Fig. 2C).

To directly observe the separation of the IDR and the OD upon condensation, we created and fluorescently labeled two cysteine variants based on the TEV cleavage construct. We opted for the TEV strategy instead of purifying each part separately, as the OD tends to aggregate during purification. We introduced a cysteine at position -20 (in the linker and HisTag region preceding the native PopZ sequence, Fig. 2A) in one construct and at position 132 (in the OD) in the other construct. We labeled them with AF647 (red) and AF488 (green), respectively. Prior to TEV cleavage, both green and red signals were diffuse. Following cleavage, the green channel showed droplets of labeled OD, whereas the red channel showed dark circles in the same regions, confirming that the OD condensate excluded H1-IDR segments (Fig. 2D). To rule out any potential dye-related effects on partitioning, we swapped the dye positions and continued to observe OD droplets and diffuse H1-IDR (Fig. S1C, top). When we formed full-length (uncleaved) PopZ droplets by adding MgCl_2_ as a control, both green- and red-labeled constructs partitioned inside (Fig. S1C, lower panels). These findings demonstrate that the OD alone is necessary and sufficient for droplet formation and that OD droplets exclude the IDR. Moreover, the condensation of the OD occurred without the addition of MgCl_2,_ low *p*H, or high salt concentrations. Thus, when covalently attached to the OD, H1-IDR inhibits condensation via a mechanism that can be overcome through low *p*H, the addition of divalent cations, or high salt concentrations.

To determine whether condensates formed by isolated OD are also composed of filaments, we inspected them using cryo-ET. In the absence of MgCl_2,_ these OD condensates (obtained through TEV cleavage) exhibited an internal structure resembling that of WT-PopZ condensates (Fig. 2E), indicating that the OD alone is sufficient for filament formation. When we analyzed the length distribution of OD filaments, we found it closely matched that of WT-PopZ, with both the median and average lengths only 3–5 nm shorter (Fig. S1D). Because this difference is smaller than the size of the building block itself (as discussed in later sections), we conclude that the IDR has minimal influence on filament length.

The mechanism of condensation through interactions between structured filaments and their inhibition via an IDR remains largely underexplored, although similar mechanisms have been documented in a few systems (Discussion). Indeed, current phase separation predictors did not capture PopZ’s condensation ability (Table S1, Fig. S2C-E), likely due to their focus on disordered domains. Thus, there is a need for predictors that can account for condensation driven by mechanisms involving structured domains, as employed by PopZ.

### PopZ forms trimers, hexamers, and multiples of hexamers

To study the assembly pathway leading to PopZ filamentation, we turned to mass photometry. This technique detects individual particles as they land on an illuminated coverslip, causing subtle changes in the laser’s scattering profile. Landing events are captured and converted into mass values based on a calibration set^39,40^. By recording thousands of such events, we generated histograms representing the distribution of the particles’ masses. When we performed mass photometry with dispersed WT-PopZ at 250 nM, it adopted trimers and hexamers (Fig. 3A, gray). The mass of the PopZ monomer (21.4 kDa) is near the lower threshold of the instrument (20 kDa) and cannot always be reliably observed.

**Figure 3.**
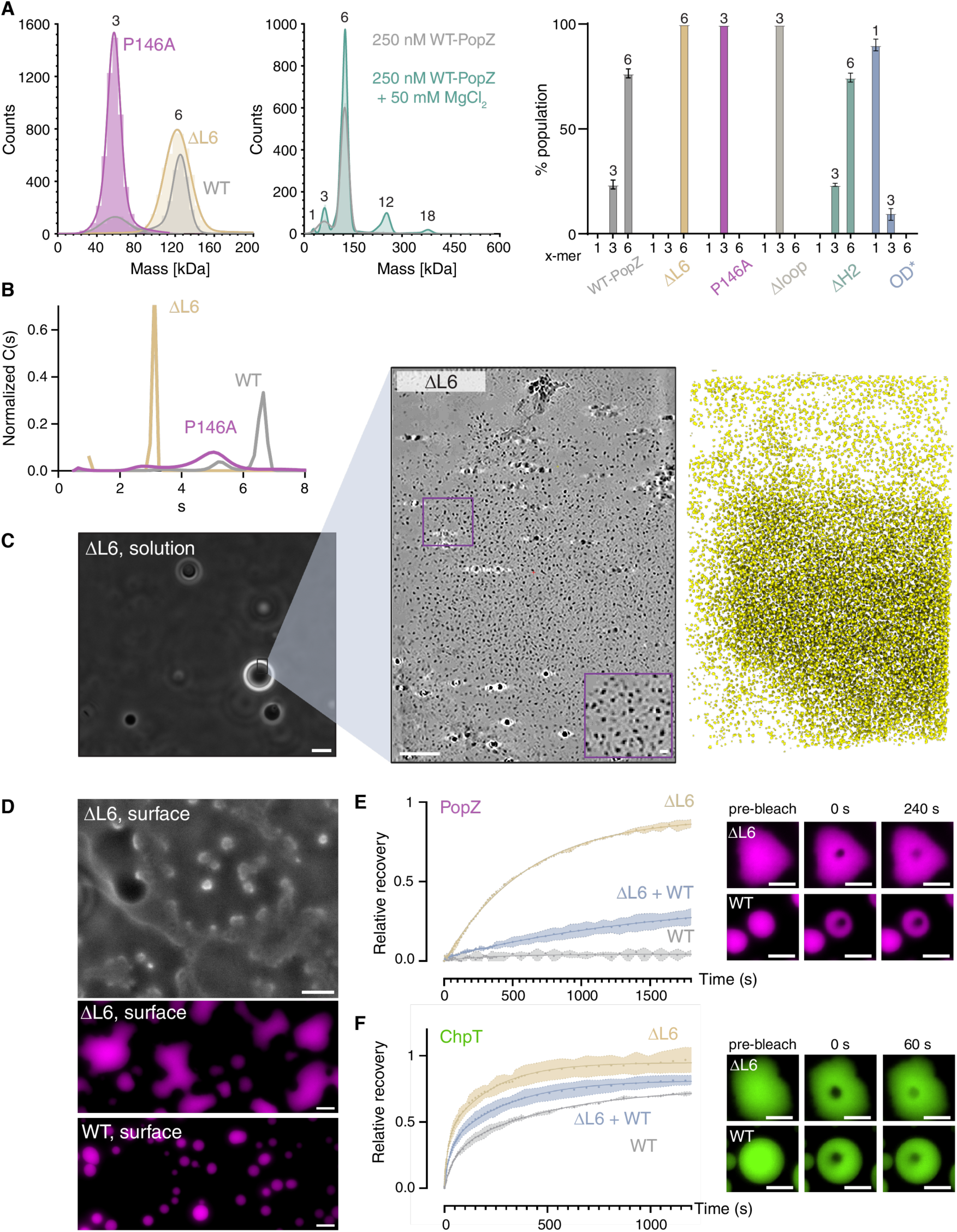
PopZ oligomerization into trimers, hexamers, and multiples of hexamers and the role of L6 in condensate formation, structure, and dynamics. **(A)** Mass photometry analysis of purified PopZ variants at 250 nM. (left) WT PopZ (gray) forms trimers and hexamers, while the P146A variant (magenta) is trapped in the trimeric state, and the ΔL6 variant (tan) is trapped in the hexameric state. (middle) Comparison of WT PopZ in the presence (turquoise) and absence (gray) of MgCl_2_ in 10 mM sodium phosphate *p*H 7.4 + 150 mM NaCl, showing that MgCl_2_ promotes higher-order oligomerization. (right) Relative populations of monomer, trimer, and hexamer across several PopZ variants, including WT (gray), ΔL6 (tan), P146A (magenta), Δloop (light gray), ΔH2 (green) and OD* (P146A-W151G-L152S, blue). Columns represent the mean of biological duplicates, and error bars represent the standard deviation. **(B)** Analytical ultracentrifugation of PopZ variants at approximately 100 µM in 10 mM sodium phosphate *p*H 7.4 + 150 mM NaCl shows that ΔL6 (tan) remains hexameric even at high concentrations, whereas P146A (magenta) and WT (gray) form higher-order oligomers, with WT assembling into the largest species. **(C)** ΔL6-PopZ forms condensates made up of hexamers. (left) Phase contrast image of ΔL6-PopZ condensates in solution. Scale bar: 5 µm. (middle) Cryo-ET tomogram of a ΔL6-PopZ condensate formed by adding 50 mM MgCl_2_ to 50 µM of purified ΔL6-PopZ in 10 mM sodium phosphate *p*H 7.4 + 150 mM NaCl. Scale bars: 100 and 10 nm. (right) 3D rendering of the traced ΔL6-PopZ tomograms. **(D)** Phase contrast (top) and fluorescence microscopy (middle) show that ΔL6-PopZ condensates wet the surface of glass coverslips, whereas WT PopZ condensates (bottom) largely remain spherical over a period of four hours. Scale bars: 5 µm. **(E)** (left) Fluorescence recovery after photobleaching (FRAP) experiments showing the recovery profiles of AF647-labeled PopZ for different condensate compositions. Mean ± standard deviation is shown for two biological replicates, each averaged over at least nine condensates. The solid lines indicate fits to a simple exponential model. (right) Representative examples of droplets used for FRAP experiments. Scale bars: 5 µm. **(F)** Deletion of the L6 region impacts client protein (ChpT) diffusivity inside the condensate. (left) FRAP recovery curves of ChpT inside WT-PopZ condensates (gray) and ΔL6 condensates (tan). Mean ± standard deviation over two biological replicates, each averaged over at least twelve condensates. The solid lines indicate fits to a two-phase exponential model. (right) Representative examples of droplets used for FRAP experiments. Scale bars: 5 µm.

To test how buffer conditions affect PopZ oligomerization, we repeated mass photometry measurements in sodium phosphate, Tris-HCl, and HEPES buffers at varying salt concentrations. While the relative populations of trimers and hexamers varied with buffer and salt concentration, both species were consistently observed above nanomolar concentrations (Fig. S2A). Because MgCl_2_ triggers condensation above the saturation concentration (*c*_sat,_ approximately 1 µM), we asked whether it affects PopZ oligomerization at lower concentrations and repeated mass photometry with 250 nM PopZ and 50 mM MgCl_2_. Although PopZ concentration was below *c*_sat_, MgCl_2_ indeed promoted the formation of higher-order oligomers, specifically 12-mers and 18-mers (Fig. 3A, middle panel), suggesting that PopZ stacks hexamers but not trimers. Thus, we propose that these hexamers serve as building blocks for PopZ filaments. Consistent with this interpretation, low *p*H and increased salt concentrations also promoted hexamer stacking (Fig. S2B). These conditions likely screen the repulsive negative charges of the IDR and thereby allow ODs of different hexamers to approach each other and interact.

To further elucidate the determinants of oligomeric assembly, we examined two PopZ variants that were shown to disrupt oligomerization in native gel electrophoresis and multi-angle light scattering^35^: P146A and Δ172–177 (hereafter referred to as ΔL6, indicating the deletion of the last six residues). At 250 nM, P146A formed only trimers, whereas ΔL6 assembled exclusively into hexamers (Fig. 3A). The addition of MgCl₂ did not induce higher-order oligomerization in either variant (Fig. S2A, right panel); P146A remained trimeric, and ΔL6 remained hexameric. Surprisingly, using light microscopy, we observed that both variants still formed condensates, albeit at higher *c*_sat_ than the WT—approximately 2 µM for P146A and 20 µM for ΔL6 versus <1 µM for WT-PopZ (Table S2). We therefore sought to further interrogate the relationship between filamentation and condensate formation.

Mass photometry is unsuitable for monitoring oligomerization at micromolar concentrations, as it becomes challenging to distinguish between a single large particle landing and two smaller particles landing close to each other. To study the oligomerization of dispersed PopZ in the micromolar range, we turned to analytical ultracentrifugation (AUC)^41^. AUC analysis revealed distinct oligomerization patterns for P146A, ΔL6, and WT-PopZ (Fig. 3B). The WT protein (gray) produced the largest species, which we attribute to filaments of varying lengths. P146A (magenta) also formed higher-order oligomers, but these species were smaller than those observed with WT-PopZ. ΔL6 (tan) did not grow beyond a sedimentation coefficient of 3, which we attribute to the hexamer. Unexpectedly, WT-PopZ populated a single species with a large s value rather than a distribution similar to that observed within the condensate by cryo-ET (Fig. 1B). This might be due to the much higher concentrations required by AUC (100 µM) compared to cryo-ET and physiological conditions (5 µM), possibly leading to the formation of supramolecular clusters that limit filament growth. Because all three variants could form condensates, these data suggest that either filamentation is not strictly required for condensation or that ΔL6 requires magnesium, salt, or low *p*H to form filaments.

### Condensates devoid of filaments have altered material properties

To test whether magnesium-triggered condensation allows ΔL6-PopZ to stack hexamers into filaments, we determined the 3D structure of the ΔL6-PopZ condensate using cryo-ET. Unlike the filamentous WT-PopZ and OD droplets, ΔL6-PopZ droplets were devoid of filaments (Fig. 3C). In addition, these droplets exhibited extensive wetting of the glass surface (Fig. 3D). This observation suggests that while ΔL6-PopZ can still condense without forming filaments, the reduced multivalency of hexamers compared to filaments likely leads to the increase *c*_sat_. The loss of filaments also impacted the overall shape of the condensates. The tendency of a liquid to minimize its surface area by forming a sphere is related to surface tension, an important material property in condensate biology^14,42^. WT-PopZ droplets (Fig. 1A) appeared spherical with clear boundaries, indicative of high surface tension. In contrast, ΔL6 droplets had irregular boundaries and deformed upon contact with the glass surface, displaying enhanced wetting behavior (Fig. 3D and Fig. S3 for a detailed comparison of WT, OD, and ΔL6). These observations indicate that loss of filamentation drastically affected the surface tension of PopZ condensates, highlighting the direct relationship between condensate substructure and emergent properties.

Another biologically important material property is viscosity, which is related to the diffusivity of molecules within a condensate. Diffusivity is often probed via fluorescence recovery after photobleaching (FRAP^43^). To test if filamentation affects the diffusivity of PopZ, we incubated fluorescently labeled full-length PopZ with either WT, ΔL6, or a mix of both and performed FRAP experiments (Table S3 for fitting results). In condensates made up of exclusively WT-PopZ (*i.e.,* filaments), virtually no recovery was observed after 20 minutes. When WT-PopZ was supplemented with hexameric ΔL6, we observed some recovery and could obtain mobile fraction and half-time of recovery (40% and approximately 27 min). This effect was amplified in condensates made up exclusively of ΔL6 (Fig. 3E, upper right panel; 90% mobile fraction and 10 min recovery half-time). Thus, PopZ could move more freely through condensates made up of hexamers compared to filaments. As with surface tension, these results provide a structural basis for condensate emergent properties.

Next, we addressed whether the loss of filaments also affects the diffusivity of condensate clients. We define clients here as components that partition into and reside within pre-formed condensates^44^. In *C. crescentus*, PopZ interacts with clients via its H1^37,45^, which is distant in sequence from the OD and should be unaffected by the deletion of the last six residues. We examined the recruitment of the client ChpT^46^, an essential mediator within *Caulobacter*’s cell-cycle circuit that directly binds to PopZ^29^. We labeled ChpT with AF488 and observed its recovery after photobleaching in WT, mixed, and ΔL6 droplets (Fig. 3F, lower right panels). The recovery kinetics for all FRAP experiments with ChpT required a two-phase model, which includes a fast and slow component, indicating that more complex dynamics contribute to the recovery, likely a combination of diffusion and binding/unbinding between ChpT and PopZ. Increasing the hexamer fraction in the condensates (and thus decreasing the filament fraction) again sped up recovery in both the fast and slow components and increased the mobile fraction (Table S3). These results indicate that filamentous WT sequesters ChpT more effectively than the hexameric ΔL6-PopZ. In summary, the last six residues (172–177) contribute to the formation of filaments, which in turn affect the condensate’s material properties. Because WT-PopZ can form filaments without condensing and ΔL6-PopZ can condense without forming filaments, we conclude that filamentation and condensation are two distinct processes governed by different residues or driving forces.

### The conserved H3H4 helices form a trimeric coiled coil

Above, we have shown that PopZ forms trimers, hexamers, and filaments. A structural description of these oligomeric states and how they interact to form condensates will provide deeper insights into PopZ biology and facilitate targeted sequence modifications for engineering purposes. To identify the determinants of PopZ oligomerization, we combined mass photometry and mutagenesis with structural models computed by AlphaFold 2 (AF2)^47^ and AlphaFold 3 (AF3)^48^. AF2 confidently predicted that the trimeric form of PopZ features a hydrophobic core formed by a coiled coil of three H3H4 helices (residues 136−177, Fig. S4A) and stabilized by the hydrophobic effect. This core remained stable when the H2 helix (residues 102−128) and the flexible linker connecting H2 to H3H4 (residues 129–135, which we refer to as “the loop”) were included in the model. In these predictions, the three H2 helices adopted variable orientations relative to H3H4, likely due to the flexibility of the loop connecting these segments (Fig. S5A). To experimentally validate these predictions, we attempted to destabilize the putative hydrophobic core by mutating W151 and L152 (highlighted in Fig. S4A) to glycine and serine, respectively, in the background of the P146A variant. During purification, this construct was prone to aggregation, and mass photometry confirmed that it existed primarily as a monomer (Fig. 3A, right panel, OD* in blue), confirming that W151 and L152 are indeed crucial for the stability of the trimeric core. Given the significance of this trimeric core, we hypothesized that it is a conserved feature across PopZ orthologs in α-proteobacteria. Using AF2, we predicted the structures of PopZ trimers from orthologs spanning hundreds of species across α-proteobacteria (Khan et al., in preparation). In all cases, the predicted structures closely resembled the trimeric coiled coil of *Caulobacter* PopZ (Fig. S4A), suggesting that the trimeric core is a central feature of PopZ.

### Hexamers are conformationally heterogeneous and stabilized by the loop

To characterize PopZ hexamers, we employed an integrative approach, combining AF2 and AF3 structural models with experimental validation. We tested the models through mutagenesis and mass photometry to determine oligomeric states and cryo-electron microscopy (cryo-EM) techniques—including sub-tomogram averaging and single-particle cryo-EM 2D class averages—to resolve the hexamer shape and dimensions. This approach allowed us to identify the sequence regions required for hexamer formation.

We began by truncating the PopZ sequence around the trimeric H3H4 core to identify a reduced segment required for hexamer formation. First, we designed a construct missing the last six residues with a TEV cleavage site between the IDR and the OD (H1-IDR-TEV-OD-Δ6) and purified it. Following cleavage, we separated H1-IDR from OD-ΔL6 using ion exchange chromatography (Fig. S5B). Isolated OD-ΔL6 (residues 102−171) remained soluble at concentrations below 5 µM but formed droplets at higher concentrations, with mass photometry confirming hexamer formation (Fig. S5B, right panel), indicating that neither L6 nor H1-IDR is necessary for hexamer assembly.

Next, we used mass photometry to examine ΔH2-PopZ, a variant lacking residues 102–128, and observed trimer and hexamer populations comparable to those of full-length PopZ (Fig. 3A, right panel, gray and green), suggesting that H2 is also dispensable for hexamer formation. Finally, mass photometry of Δloop-PopZ, lacking residues 129–134, showed a complete shift of the population to trimers (Fig. 3A, right panel, light gray), indicating that the loop connecting H2 to H3H34 is essential for hexamer formation (see Fig. 6A for a summary of sequence determinants). Taken together, these mass photometry experiments with truncated PopZ variants identified residues 129−171 as a reduced hexamer-forming segment.

To define the hexamer shape and dimensions, we performed sub-tomogram averaging on hexameric ΔL6-PopZ. Given that cryo-ET typically does not resolve disordered segments due to their conformational variability and lack of defined structure, the class averages we obtained can be attributed to the structured OD. Our sub-tomogram averaging analysis resolved two predominant hexamer shapes (Fig. S4B): one straight, measuring 100 Å in length, and the other curved, extending 130 Å.

We used AF2 to model the PopZ hexamer with the reduced hexamer-forming segment, consisting of six chains of residues 129–171. The model revealed head-to-head interactions between trimers mediated by the loop and H3 helices, with C-termini protruding outward (Fig. S6A). These features are in agreement with our subtomogram averaging of ΔL6-PopZ (Fig. S4B) and our findings that hexamer formation requires the loop but not the last six residues nor H2. The model also suggests that multiple coiled coil registers are accessible, as indicated by the sliding between H3H4 helices when compared to the trimer model (Fig. S6A, lower left panel). Additional support for the AF2 model came from single particle cryo-EM 2D class averages of the ΔH2ΔL6-PopZ construct (Fig. S6B), though the complex’s inherent flexibility limited high-resolution structural determination. We also computed a hexamer model including H2 (residues 102−128), where a bundle of H2 helices replaced the H3 contacts, resulting in an S-shaped conformation compatible with the curved 2D classes obtained in subtomogram averaging of ΔL6-PopZ (Fig. S6C). Together, these results indicate that H3H4 helices form trimers that interact through the loop to assemble hexamers approximately 100– 130 Å in length. Furthermore, the model implies that the hydrophobic H2 helices project outward from the hexamer center, potentially forming interoligomer contacts with H2 bundles from other PopZ particles.

### PopZ filaments grow longitudinally from hexameric building blocks

To gain high-resolution insights into WT-PopZ filaments, we employed subtomogram averaging on the segmented WT-PopZ filaments. This approach proved challenging due to the heterogeneity and pronounced curvature of the filaments (Fig. 1A, lower panels). To address this, we focused our analysis on a subset of straight filaments, which we isolated using a segmentation-based particle picking method. The class averages revealed that the filaments have the same width as individual hexamers, suggesting that filaments are formed through the longitudinal stacking of hexameric subunits (Fig. S4C). Given that H2 is not required for hexamer formation (Fig. 3A), we explored its potential role in filament assembly. AUC showed that ΔH2-PopZ did not sediment beyond a coefficient of approximately 3, indicating that it remained hexameric, similar to ΔL6-PopZ (Fig. S4D). This finding suggests that H2 contributes to filamentation either directly or indirectly.

To identify the specific residues within H2 involved in filament formation, we generated partial truncations of H2, deleting either residues 102−109 or 110−122. Surprisingly, neither construct restored filamentation (Fig. S4D, coral and magenta), indicating that the entire H2 is necessary for this process. Therefore, both the H2 region and the last six residues are required for filamentation, either because they directly participate in the filament-forming interactions or because they enable PopZ to adopt a conformation needed for filamentation. The terminal position of the last six residues makes them more likely to directly facilitate the longitudinal stacking of hexamers, while the central position of H2 suggests it may stabilize a filament-favoring conformation. Taken together, our data indicate that PopZ filaments form through longitudinal stacking of hexamers, can bend considerably, and require both the last six residues and H2 for proper assembly.

### Filaments form contacts mainly through H2

To undergo condensation, intermolecular interactions must become thermodynamically more favorable than intramolecular interactions and interactions with the solvent^49^. Because PopZ forms supramolecular assemblies (*i.e.,* oligomers and filaments), the term intermolecular can be ambiguous. Therefore, we use the term interoligomer contacts to describe interactions between distinct PopZ particles, whether they are hexamers or filaments. To identify the regions responsible for interoligomer contacts, we performed a series of deletions along the PopZ sequence and assessed their impact on droplet formation using light microscopy.

As noted earlier, deleting the last six residues abolished filament formation and significantly increased *c*_sat_ (Table S2). This increase likely stems from a reduction in multivalency due to loss of filaments without compromising interoligomer contacts. Similarly, deletion of the loop also led to an increase in *c*_sat_, though condensation still occurred. In contrast, deleting the H2 region (102−128) completely prevented condensation in our standard buffer (10 mM sodium phosphate *p*H 7.4, 150 mM NaCl, 50 mM MgCl_2_), even at high protein concentration (100 µM). This result indicates that H2 contributes to contacts between PopZ particles. Although ΔH2-PopZ formed droplets at elevated salt or magnesium concentrations (Table S2), these droplets dissolved after ∼20 minutes, suggesting that while other motifs within PopZ may contribute to interoligomer contacts, H2 plays a key role in stabilizing these interactions.

To determine whether specific parts of the H2 region could restore condensation under standard conditions, we repeated light microscopy experiments with Δ102−109 and Δ110−122. Neither deletion restored condensation, possibly due to destabilization of the H2 helix. These results suggest that H2 may promote condensation, not only through specific residues but also by maintaining its helical structure. In summary, while deletions within the OD that disrupt oligomerization or filamentation also increased *c*_sat_, removal of H2 precluded condensation under standard conditions altogether. Additionally, the solvent-exposed surfaces of H3H4 may contribute to interoligomer contacts, although directly testing this hypothesis without disrupting the trimeric core (Fig. 6A) remains challenging.

As an orthogonal approach to identify the contacts between trimers, hexamers, and/or filaments that drive condensation, we conducted coarse-grained molecular dynamics simulations^50,51^ of PopZ condensation. We modeled PopZ condensates at residue resolution using the OpenABC implementation of the hydrophobicity scale (HPS) model with the Urry hydrophobicity scale^52,53^. To simulate coexistence of the dense and dilute phases, we combined this HPS model with slab simulations^54^, placing 100 PopZ trimers in a simulation box (Methods). We opted to simulate the condensation of trimers instead of hexamers due to the greater confidence in our trimer model. The trimer interface (residues 136−171) was stabilized with structure-based potentials^55^, whereas other inter-chain interactions between PopZ monomers were scored according to the force field.

First, to increase our confidence in the simulation results, we assessed whether the simulations could reproduce key experimental findings, including the conditions that promote PopZ condensation and the effect of removing H1-IDR. The final frame of the slab simulations for each of the four different conditions was consistent with our experimental observations (Fig. S4E). WT-PopZ did not form condensates at neutral *p*H and low salt concentrations. Lowering the *p*H or adding salt drove the formation of large clusters, indicative of condensation. Finally, simulations ran without H1-IDR produced a stable condensate, with only minimal amounts of trimers in the dilute phase.

With the validation of simulations in hand, we proceeded to identify the residues that drive the interactions between PopZ particles in the condensates. We examined the inter-trimer contact map of WT-PopZ during cluster formation induced by 600 mM monovalent salt (Fig. S4E, right). We identified interaction hotspots involving residues within H2, the middle of the H3H4 helix, and the loop connecting these regions. Thus, the simulations indicate that H2 and H3H4 drive trimer:trimer interactions. It is likely that similar interactions among trimers, hexamers, and/or filaments drive condensation *in vivo*. In summary, both experimental and simulation data suggest that residues across H2, the loop, and the middle of H3H4 can form interoligomer contacts, with the largest contribution coming from H2.

### H1 interacts with the OD

We have shown above that the H1-IDR segment inhibits OD condensation unless divalent cations, high salt, or low *p*H are present (Fig. 2D). To elucidate the mechanism underlying this inhibition, we employed single-molecule Förster Resonance Energy Transfer (smFRET^56,57^).

More specifically, we monitored the proximity of the N-terminus to the OD to determine if H1-IDR behaves simply as a disordered chain or if it can interact with other parts of the protein. We labeled the same cysteine positions as in Fig. 2D (C-20 and C132) within the same molecule. We used mass photometry and light microscopy to verify that the protein was still able to form oligomers and condensates after introduction and labeling of cysteine (Fig. S7A). For smFRET experiments, we supplemented this PopZ FRET reporter (at ∼300 pM) with a 16,667-fold excess of unlabeled WT-PopZ (5 µM) to prevent formation of oligomers with more than one FRET reporter, which could lead to artifacts in FRET parameters.

Based on the distance between the labeled cysteines, we expected a low FRET efficiency. With 151 disordered residues between the donor dye (AF488) and the acceptor dye (AF647), we anticipated a mean distance of about 91 Å for a random chain^58^, which corresponds to a FRET efficiency (E_FRET_) < 0.1. Because the IDR of PopZ is enriched in negative charges and proline residues, we also employed the deep-learning algorithm ALBATROSS, which predicts end-to-end distances of IDRs based on their sequence attributes^59,60^. ALBATROSS predicted end-to-end distances exceeding 100 Å (corresponding to E_FRET_ < 0.1) for both the full-length sequence of our FRET reporter and the region between the two dyes.

We indeed observed a population at low E_FRET_ ∼0.1 (Fig. 4A). However, we also observed an intermediate E_FRET_ state (mean E_FRET_ > 0.5, average inter-dye distance of ∼54 Å), suggesting that the N-terminus and the OD can come much closer to one another than expected in the absence of intra-PopZ interactions. Using photon distribution analysis^61^ we found that the fraction of PopZ adopting the compacted (intermediate E_FRET_) state is approximately 67%, or two-thirds of the total population. To increase our confidence in this result, we used 6 M Gdn-HCl to denature PopZ and weaken intermolecular interactions, which led to a significant decrease in the compacted state (Fig. 4B, right panel, tan).

**Figure 4.**
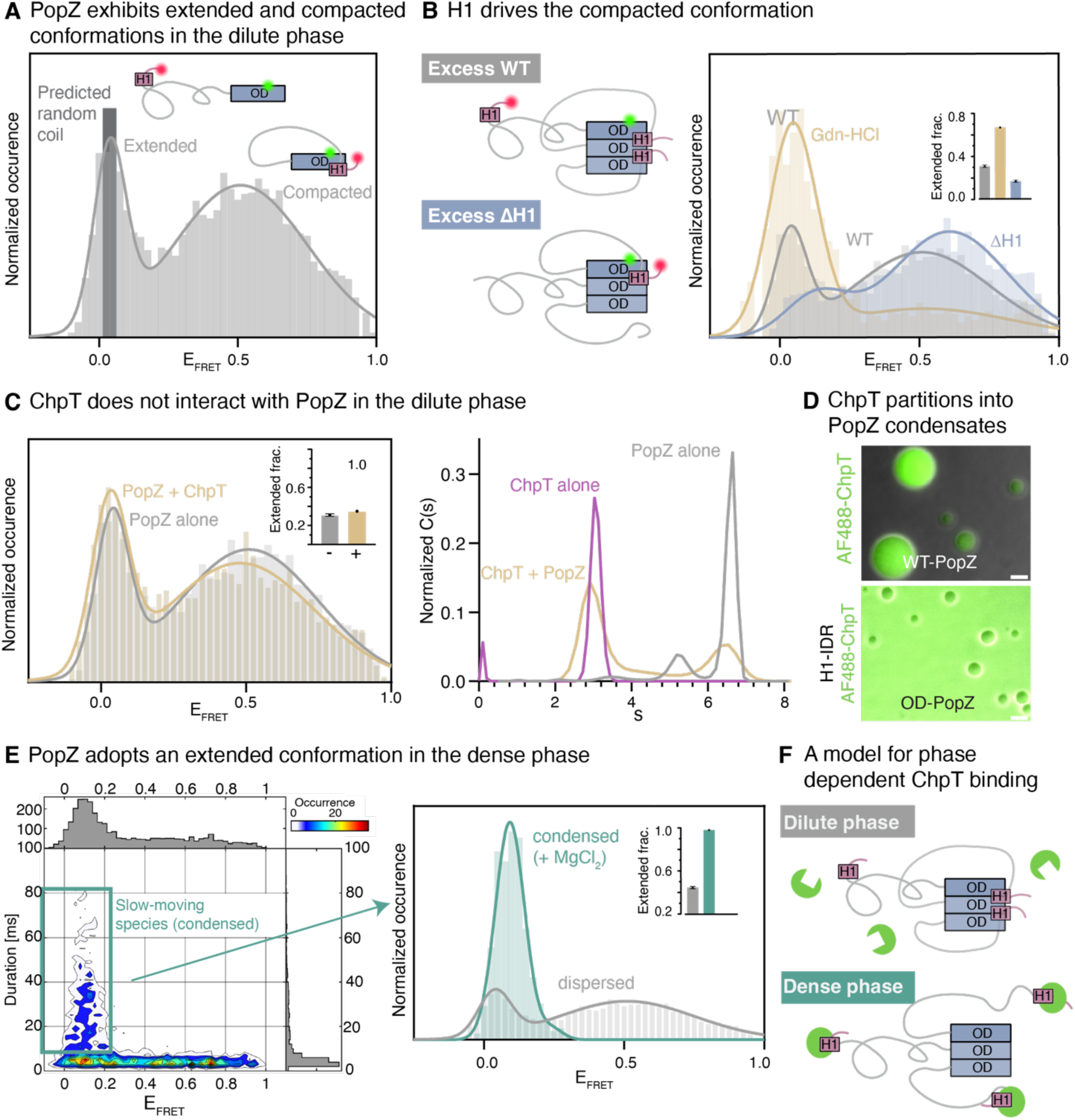
The conformational equilibrium of PopZ is phase-dependent. **(A)** smFRET reveals that PopZ adopts extended and compacted conformations in the dilute phase. FRET efficiency distribution of the smFRET reporter labeled at C-20 and C132 shows two predominant conformations in the dilute phase (light gray): extended and compacted. The extended conformation coincides with the predicted behavior of a random coil (dark gray), whereas the compacted conformation indicates proximity between the N- and C-termini. **(B)** H1 drives the compacted conformation. (left) Schematic illustrating two experimental conditions. In the first condition, labeled FL-PopZ was mixed with an excess of unlabeled WT PopZ. For each labeled PopZ monomer (fluorophores indicated by green and red circles), two unlabeled PopZ monomers are shown, where the H1 of the unlabeled monomer competes with the H1 of the labeled monomer for a binding site on the OD (excess WT, top). In the second condition, labeled FL-PopZ was mixed with an excess of unlabeled ΔH1-PopZ (missing residues 1-23). In this case, only the H1 of the labeled PopZ monomer can bind the OD (excess ΔH1, bottom). (right) FRET efficiency plot shows that excess WT promotes the extended conformation (gray), similar to the denaturing condition (tan). In contrast, excess ΔH1-PopZ promotes the compacted conformation (blue). **(C)** H1 recruits ChpT to PopZ condensates. Fluorescence microscopy images show 1 nM ChpT labeled with AF488 recruited by FL-PopZ condensates (top), while ChpT remains diffuse in the presence of condensates of isolated OD (bottom). In both conditions, PopZ concentration was 5 μM. Scale bar: 5 µm. **(D)** ChpT does not interact with PopZ in the dilute phase. (left) FRET efficiency histograms indicate that ChpT at 5 µM (tan) only marginally increased the population of the extended conformation of PopZ (gray). (right) Sedimentation coefficient distribution from AUC for individual samples of PopZ (gray) and ChpT (magenta) as well as combined (tan). The combination did not indicate the formation of a new PopZ:ChpT species. **(E)** In the dense phase, PopZ adopts exclusively the extended conformation. (left) 2D histograms of FRET efficiency versus burst duration. Typical burst durations for single protein particles are shorter than 10 ms. Durations greater than 15 ms (turquoise box) were selected to isolate fluorescence bursts coming from within condensates. (right) Condensed PopZ exclusively adopts the extended conformation. FRET efficiency histogram of condensed and dispersed (same condition as in (A)) PopZ. **(F)** A model of phase-dependent interactions. In the dilute phase, H1 is predominantly bound to the OD, preventing ChpT (green) from interacting with H1. In the dense phase, H1 is released from the OD, enabling ChpT binding.

Because the only structured element in the N-terminus is the H1 helix, we hypothesized that H1 binds the OD. If this interaction occurs, not only would H1 of the FRET reporter we are monitoring be able to bind the OD, but so would the H1 helices of the unlabeled PopZ molecules. To reduce competition between H1 of labeled and unlabeled PopZ, we supplemented the FRET reporter with unlabeled ΔH1-PopZ instead of WT (illustrated in Fig. 4B, left panel). This significantly increased the population of the compacted state, confirming that it is H1 that mediates the interaction leading to compaction of PopZ (Fig. 4B, right panel, blue). This interaction is also predicted by AF3, although the pLDDT and the overall model scores are low (Fig. S7B).

### H1 suppresses interfilament interaction in the dilute phase and exposes client-binding sites in the dense phase

Given that H1 is responsible for client binding, we expected that clients like ChpT would bind to and therefore stabilize the extended state of PopZ. To our surprise, the addition of 5 µM ChpT had almost no effect on the extended population (Fig. 4C), although this ChpT concentration should have been saturating based on previous studies of the ChpT:PopZ interaction^29,37^. These two studies, however, may have altered PopZ behavior—Lasker et al. immobilized PopZ on an SPR cell surface, possibly affecting its fold and oligomerization, while Holmes et al. removed the OD to make PopZ amenable for NMR, thereby eliminating the compacted state entirely. To test whether ChpT binds to dispersed PopZ with orthogonal techniques that don’t compromise its structural integrity, we turned to mass photometry and AUC. Neither technique—mass photometry analysis at 250 nM ChpT and PopZ, or AUC at 40 µM ChpT and PopZ, a concentration significantly exceeding physiological levels—revealed any detectable interaction (Fig. 4C, right panel). In contrast, condensed PopZ robustly recruited AF488-labeled ChpT (Fig. 4D), even at 100-fold lower concentrations. Recruitment was lost when we formed droplets of isolated OD, demonstrating that recruitment depends on the presence of H1-IDR. These results suggest that condensation significantly enhances ChpT:PopZ interactions.

To rationalize the lack of robust ChpT:PopZ interactions in the dilute phase, we hypothesized that the H1:OD interactions prevent H1:ChpT interactions. By implication, the robust recruitment of ChpT into PopZ condensates suggests that H1 in condensed PopZ no longer interacts with the OD. To determine whether condensation affects H1:OD interactions, we repeated the smFRET experiment on condensed WT-PopZ by adding 50 mM MgCl_2_. We plotted the duration of fluorescence bursts (the time a labeled molecule spent in the detection volume) versus E_FRET_ to select only bursts belonging to slow-moving species (*i.e.,* droplets). Fig. 4E shows that condensation shifted the entire population to the extended state, where H1 and OD are far from each other. Together, these data suggest the following model: in the dilute phase, H1 interacts with the OD, serving as a competitive inhibitor of client binding. Upon condensation, however, these H1 interactions are suppressed, relieving the inhibition and allowing the recruitment of ChpT and potentially other client partners. This mechanism combines avidity with a condensation-dependent conformational switch.

Our model implies that removing H1 would promote the extended state and might allow condensation without divalent cations, salt, or low *p*H. When we purified ΔH1-PopZ, however, we observed no formation of spherical droplets. Upon addition of magnesium, ΔH1-PopZ formed peculiar aggregate-like clusters (Fig. S7C, right panel), similar to those at low *p*H without salt or magnesium (Fig. S1B). In AUC, ΔH1-PopZ formed non-native species that grew much larger than WT-PopZ (Fig. S7C, left panel). We propose that without the tethering of H1 to the OD, PopZ can get trapped in aggregation-prone intermediates. The IDR likely plays an important role in the formation of these intermediates, as isolated OD, which lacks both H1 and the IDR, is well-behaved *in vitro* (Fig. 2).

### Filamentation is crucial for PopZ function *in vivo*

To assess the importance of PopZ filament formation *in vivo*, we expressed filament-deficient mutants in *Caulobacter* cells. Replacing WT-PopZ, which forms polar foci, with hexameric PopZ (ΔL6-PopZ) has been shown to lead to uniform cytoplasmic distribution of PopZ^35^, indicating that filaments are important for condensation—likely by increasing multivalency, consistent with our *in vitro* data. To determine whether filamentation confers functional features beyond increased multivalency, we expressed low levels of mCherry-ΔL6-PopZ in otherwise wild type cells, mimicking our *in vitro* mixing experiments (Fig. 3E-F, blue FRAP curves). Expression of low amounts of mCherry-ΔL6-PopZ, via the leaky expression observed in PYE media^35^, was toxic, inhibiting cell growth altogether (Fig. 5A).

**Figure 5.**
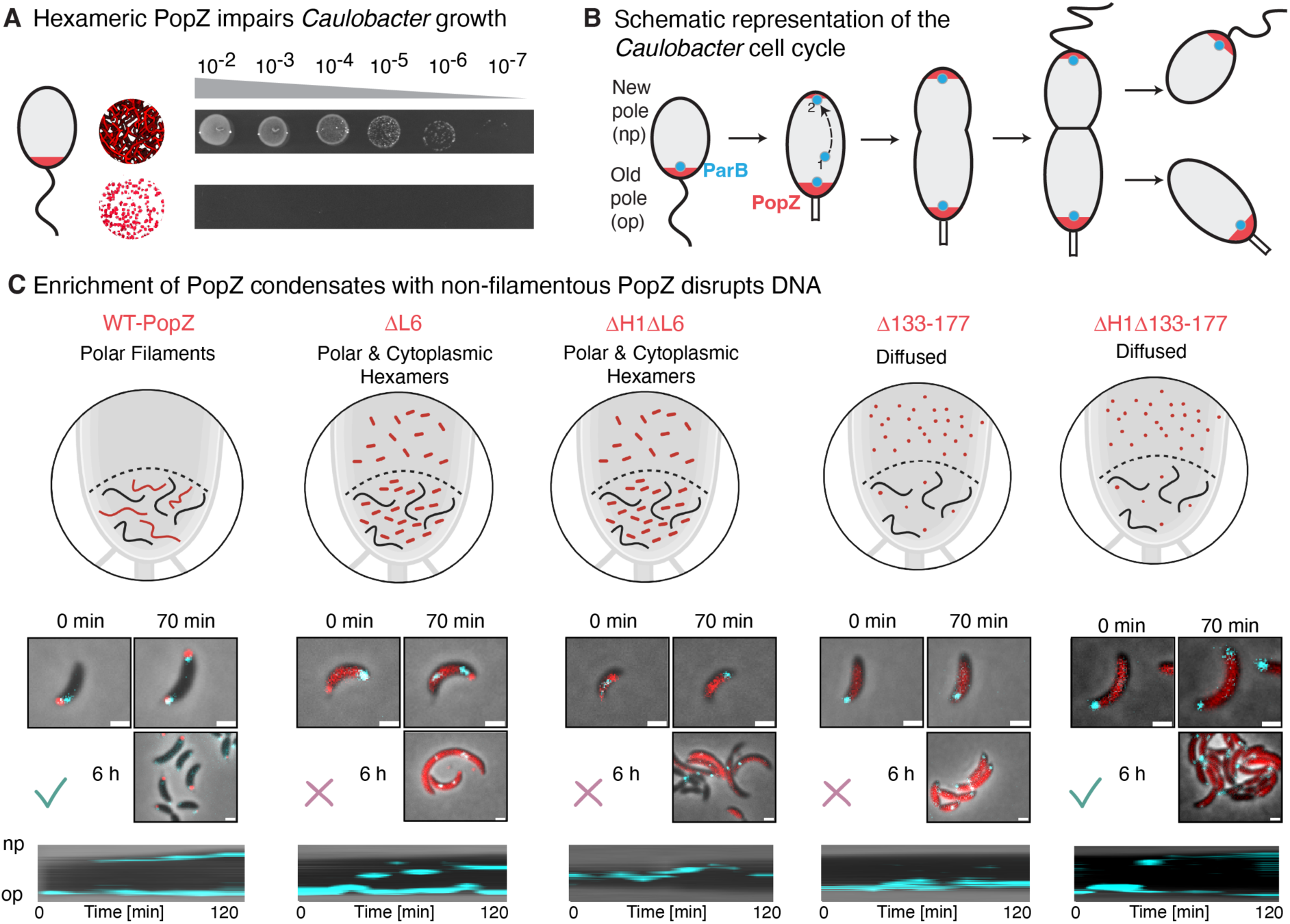
Filaments are crucial for PopZ function in vivo. **(A)** Spotting dilution assay of *Caulobacter* cells expressing mCherry-PopZ or mCherry-ΔL6-PopZ from a plasmid in a NA1000 *parB*::*cfp*-*parB* background. Cells expressing mCherry-PopZ exhibit normal growth (top), whereas cells expressing mCherry-ΔL6-PopZ (bottom) do not grow. **(B)** Schematic illustrating the migration and capture of the ParB:*parS* focus (blue) by the PopZ microdomain (red). At the start of the cell cycle, ParB forms a complex with the *parS* centromere that is anchored to the PopZ condensate at the old pole. During chromosome replication, one copy of the ParB/parS complex remains anchored at the old pole via ParB interactions, while the other copy migrates to the new pole, associating with PopZ with assistance from a ParA gradient (not shown) to facilitate rapid migration^64,65^. **(C)** The effect of PopZ mutants on DNA segregation. (top) Schematic illustrating the oligomerization pattern and localization of different mCherry-PopZ variants (red) expressed in WT *Caulobacter* cells expressing CFP-labeled ParB and endogenous WT PopZ (black). (middle) Representative time-lapse images of synchronized cells at three time points, starting from swarmer cells. The cells express mCherry-PopZ variants from a plasmid, while CFP-ParB (blue) replaces WT ParB to track DNA segregation. Check marks or X-marks indicate whether chromosome segregation proceeded properly, *i.e.,* whether ParB foci were anchored and the old pole, replicated, migrated to the new pole, and captured at the new pole within expected time frames. (bottom) Kymographs displaying the cell body (black) and CFP-ParB (cyan) over time. Time progresses from the left to the right. The old pole is positioned at the bottom, while the new pole is at the top. Scale bar: 1 µm.

To elucidate the mechanism underlying this toxicity, we monitored DNA segregation using the ParABS partitioning system^62,63^ (Fig. 5B) upon expression of different PopZ mutants. By labeling the centromere-binding protein ParB with CFP and tracking its motion throughout the cell cycle, we obtained a sensitive probe for PopZ’s ability to anchor and capture ParB foci. Expression of low levels of mCherry-WT-PopZ in addition to the native PopZ did not adversely affect ParB anchoring at the old pole, migration, or capture at the new pole. In contrast, low expression levels of mCherry-ΔL6-PopZ led to severe phenotypes. mCherry-ΔL6-PopZ was partially localized at the pole and dispersed throughout the cytosol (Fig. 5C). It also displayed increased dynamics, frequently exchanging between the pole and cytosol. Finally, the presence of hexameric PopZ significantly impaired the ability to capture and anchor ParB:*parS* complexes at the poles (kymographs in Fig. 5C).

To assess whether the observed aberrations in ParB:*parS* anchoring were due to impaired PopZ structure at the pole, its presence in the cytosol, or both, we examined two additional mutants: Δ133–177, lacking the H3H4 region and incapable of condensation and polar localization and ΔH1Δ133–177, which is unable to bind clients (Fig. 5C). While Δ133–177 drastically slowed down ParB motions, deletion of H1 restored normal function, suggesting that cytoplasmic PopZ that is unable to interact with clients is not harmful. However, when we deleted H1 from ΔL6-PopZ, cells were still unable to anchor or capture ParB:*parS* foci, indicating that ΔL6-PopZ impairs *in vivo* function by interfering with the WT-PopZ condensate rather than through aberrant client interactions. In summary, our results indicate that filamentation is crucial for the function of PopZ condensates *in vivo,* not merely by enhancing multivalency but by maintaining condensate integrity essential for proper DNA segregation.

## Discussion

### A mechanistic model of PopZ condensation

The objective of this study was to characterize the relationship between the primary sequence of PopZ and its supramolecular assembly into condensates. We uncovered roles for the H1 helix and the IDR in modulating condensation and gained insight into how the OD drives oligomerization, filamentation, and condensation. Our mechanistic model (Fig. 6B) proposes that PopZ monomers form a trimeric coiled coil via H3 and H4 through hydrophobic interactions. Trimers then dimerize in a head-to-head orientation via the loop and either H2 or H3. This dimerization results in a hexamer where the C-termini of all six monomers point outwards in two directions along a central axis. The loop, H2, IDR, and H1 chains sprout outwards from the center of this hexamer.

**Figure 6.**
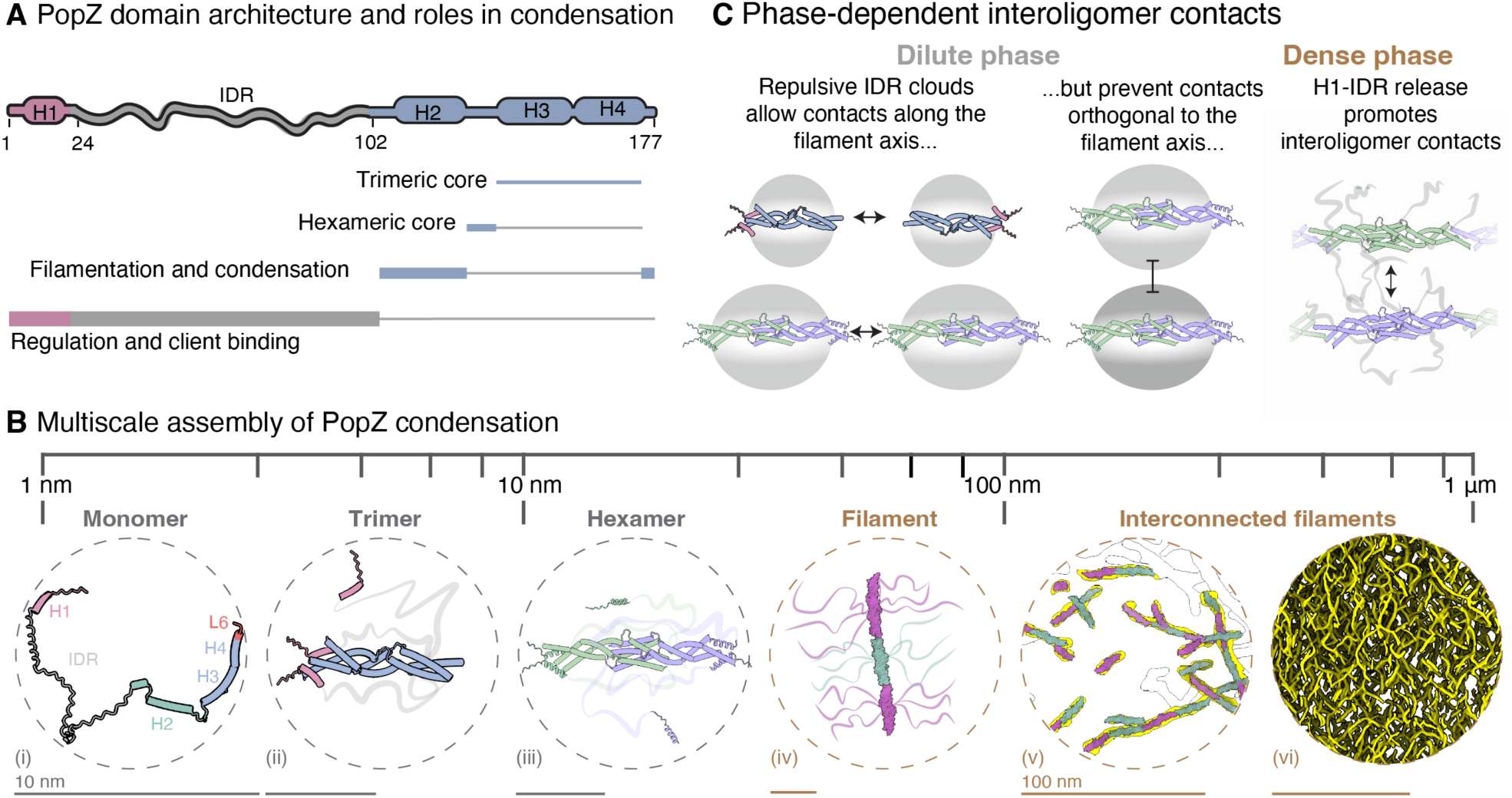
Mechanistic depiction of PopZ oligomerization, filamentation, and condensation. **(A)** Schematic representation of the PopZ sequence highlighting its subdomains and their roles in assembling PopZ into oligomers, filaments, and condensates. **(B)** Illustration of the hierarchical assembly of PopZ from monomers to condensates. (i–iii) PopZ oligomers in the dilute phase (dashed gray circles). (i) AlphaFold 2 model of the PopZ monomer, with the N-terminal domain—including the client-binding helix (H1)—in pink, the IDR in gray, the H2 helical region in green, helices three and four (H3-H4) in blue, and the C-terminal six amino acids in red. (ii) AlphaFold model of the PopZ trimer, showcasing the formation of a triple coiled coil via the H3-H4 region (blue), with H1 helices (pink) interacting proximally to these coiled regions. Conformational flexibility is illustrated by depicting two H1 helices close to H3-H4 and one further away. (iii) AlphaFold hexamer model displaying interactions between two trimers (colored purple and green) mediated by inter-trimer H2 region interactions. The IDRs extend from the center of the hexamer, creating a negatively charged cloud around it. Four of the six H1 helices are shown proximal to their corresponding H3-H4 regions. (iv-vi) PopZ filament structure in the dense phase (dashed brown circles). (iv) Fitting of three hexamers (purple and green) into a density extracted from the condensate tomogram (yellow), demonstrating a tail-to-tail organization between hexamers with IDRs extending from the center of each hexamer. In this configuration, the IDRs are positioned away from H3-H4 and are involved in binding client molecules. (v) A larger segment of the tomogram (yellow) with fitted hexamers, illustrating variability in filament lengths. Unused portions of the tomogram are outlined in black. Here, only the OD of each hexamer is displayed. (vi) Portion of a tomogram of the PopZ condensate (from Fig. 1) revealing the filamentous ultrastructure of the condensate. A scale bar is shown for each circle. **(C)** Phase-dependent interoligomer contacts. (left) Trimer and hexamer interactions in the dilute phase, with the gray ellipse representing a cloud of negative charge due to negatively charged residues and the flexibility of the IDR. (right) In the dense phase, conformational changes opening the H1 helices remove the IDR cloud around hexamers, enabling interfilament contacts.

When PopZ is dispersed, H1 can interact with the OD, looping the negatively charged IDR around it. Due to the flexibility of the IDR, we conceptualize the IDR chains as a cloud of diffuse negative charge that decreases in density with increasing distance from the hexameric core (Fig. 6C). The repulsion between these clouds of negative charge prevents PopZ particles from forming interoligomer contacts. Filamentation likely occurs longitudinally from the C-termini, where repulsion is weaker, explaining the formation of filaments in the absence of either salt or magnesium. When either salt or magnesium are added, the negative charges of the IDR are partially screened, allowing sufficient contacts between PopZ particles to culminate in the formation of condensates (Fig. 6B). Upon condensation, the H1-IDR chains do not participate in these contacts, as evidenced by the exclusion of isolated H1-IDR chains from OD condensates (Fig. 2 C-D). Instead, H1-IDR chains extend away from the OD, exposing H1 and making it more available to clients (Fig. 4F).

The functional roles of the various PopZ segments can thus be organized into a hierarchical sequence where the highly conserved H3H4 forms the trimeric core unit (Fig. 6A). Loop residues stabilize the hexamer. H2 and the L6 drive formation of filaments and interfilament contacts, leading to condensation. The IDR and H1 domains regulate condensation and material properties as well as client binding.

### Polymerization and structure

Polymerization takes center stage in the condensation behavior of PopZ. By stacking hexameric building blocks into filaments, the valency of the monomer increases manifold, allowing WT-PopZ to condense at submicromolar concentrations. Because each monomer chain contains H1, polymerization also leverages the avidity effect. That the structured OD is responsible for polymerization is not surprising, as folded proteins are generally more effective at forming long-lived oligomers than intrinsically disordered proteins (IDPs). However, polymerization and structure provide additional unique advantages in the context of condensates. The suitability of IDPs for condensate formation lies in the combination of interaction motif redundancy and flexibility, which allows multiple IDPs to readily rearrange, optimizing intermolecular interactions. Our cryo-ET data show that PopZ filaments, too, can bend and deform. Consistent with this perspective, the length distribution of PopZ filaments is heterogeneous, and many hexamers are present in condensates, implying that space can be filled by less bulky species (hexamers and short filaments) to minimize the free energy of the system. Thus, filaments formed by the structured OD retain a degree of interaction redundancy and flexibility, similar to that of IDPs, but on larger length scales. This increase in length significantly affects material properties. First, viscosity generally increases with polymer length: longer polymer chains increase drag and intermolecular interactions, and thus viscosity. Longer chains are also more prone to entangle with each other, which further increases viscosity^66,67^. Indeed, we found that WT-PopZ condensates exhibit decreased fluidity compared to ΔL6-PopZ condensates (Fig. 3E). Second, while the relationship between surface tension and polymer length is more complex^68^—especially when long side chains (*e.g.*, H1-IDR sprouting from the central filament axis) are present—longer polymer chains tend to lead to higher surface tension^69,70^.

### A phase-dependent conformational switch

We observed that H1 interacts intramolecularly with the OD in dispersed solution but not in condensates. Because H1 mediates client binding, this property, coupled with the increase in avidity accompanying condensation, creates marked differences in apparent binding affinity to clients between the condensed and dispersed phases. While further exploration is needed to fully elucidate this mechanism, it hints at promising synthetic applications. Specifically, this mechanism could enable engineering systems where target protein recruitment is activated by condensation, while cellular localization remains unaffected in the dispersed state.

### Filaments in other condensate systems

Formation of condensates requires multivalent interactions^1,71^, which have been observed in all types of biological macromolecules and involve a variety of geometries and mechanisms^72–77^. Accordingly, biological condensates sample a diverse spectrum of supramolecular architectures, ranging from highly disordered states to nearly crystalline formations^8,78,79^. Indeed, condensation through interactions between filaments such as we studied here is not unique to PopZ but has been observed in other systems, most notably p62, F-actin, and SPOP/DAXX bodies.

In the case of p62, best known as an adaptor protein of selective autophagy, filaments grow through reversible head-to-tail polymerization of its PB1 domain^17,80,81^. This filamentation is a requirement for lysosomal targeting of p62^82^. Filament length is regulated by a few clients through several mechanisms, including physical capping of the filament, while it remains unaffected by others^19,83^. Similarly, some of its binding partners are multivalent and can bridge p62 filaments, *i.e.,* form cross-links, which increases the propensity for condensation^84^. In fact, based on *in vitro* studies, p62 filaments might be unable to condense on their own, probably due to limited intrinsic interfilament contacts.

Like p62, globular actin monomers undergo head-to-tail assembly into F-actin, the filamentous form of actin^85^. In cells, both the assembly and disassembly of actin are tightly regulated by various binding proteins that nucleate, elongate, stabilize, cap, sever, and cross-link actin filaments, thereby organizing higher-order F-actin networks^86–89^. In vitro studies have demonstrated that actin cross-linking proteins can induce phase separation of F-actin into a liquid crystal phase and spindle-shaped structures^21^.

The tumor suppressor speckle-type POZ protein (SPOP) is a substrate recognition subunit of a ubiquitin ligase complex^90–92^. By recruiting substrates to the ligase complex, SPOP facilitates their ubiquitination and degradation^90^. SPOP assembles dimeric repeating units into a double-layered filament with flexible linkers and hydrophobic interfaces connecting the two layers. Stacking of the dimeric units is isodesmic, resulting in a concentration-dependent exponential distribution of filament lengths^93–95^. Each SPOP monomer also includes a substrate-binding domain, making filaments multivalent and avid for substrates. The substrates themselves are often multivalent for SPOP, which further increases avidity and allows for cross-linking of SPOP filaments. Polymerization of SPOP is important biologically as it increases the lifetime of oligomeric SPOP:substrate complexes and the enzymatic activity with respect to monomeric SPOP^96,97^. Similar to p62, substrates that cross-link SPOP filaments can drive condensation *in vitro* and co-localization with membraneless organelles *in vivo*^20^. The SPOP substrate “death domain-associated protein” (DAXX)^98^ is a particularly interesting example as it is multivalent for SPOP and itself able to self-associate. Thus, the supramolecular architecture can change drastically as a function of their molar ratio. At low DAXX/SPOP ratios, large amorphous structures form *in vitro* and are thought to be long SPOP filaments cross-linked by the few available DAXX molecules whose binding sites are saturated as a result. These structures can relax into liquid-like condensates at higher DAXX/SPOP ratios, presumably because the increased abundance of DAXX molecules leads to the formation of brush-like structures that leave free DAXX binding sites exposed. These free binding sites are thought to engage in DAXX:DAXX interactions that lead to liquid-like condensates^20,94,99^. In the first case, long SPOP filaments being cross-linked by DAXX lead to a gel-like phase. In the second case, SPOP polymerization leverages an increase in the multivalency of DAXX for itself, creating a liquid phase.

From these examples of filaments or supramolecular structures within condensates, three key mechanisms emerge: the thermodynamics of polymerization, growth capping, and cross-linking or interoligomer contacts. We use the term cross-linking to describe the bridging of two particles (*e.g.,* filaments or oligomers) via a multivalent ligand and interoligomer contacts to describe direct interactions between two particles. Because polymerization can drastically impact material properties, biological systems have evolved multiple mechanisms to regulate filament length within condensates: When filaments polymerize reversibly, their length can be kept short simply by thermodynamics (*e.g.,* isodesmic polymerization of PopZ), and a capper may not be necessary. When polymerization is cooperative or highly variable, capping molecules or substrates are needed to regulate filament length. Similarly, the strength and number of cross-links or interoligomer contacts affect the material state of the condensate and is therefore regulated either intrinsically (*e.g.,* limited by the inherently available interactions between PopZ filaments) or extrinsically (limited by the number of binding sites accessible to the substrate doing the cross-linking, or the number of available cross-linkers). While the examples above need extrinsic factors to regulate either filament length or cross-linking, PopZ has these regulatory mechanisms built into a compact helical domain, allowing it to work in isolation from other cellular systems. This idea is backed by the *in vitro* data presented here but also by foci formation and client recruitment of PopZ in orthogonal systems such as human and *E. coli* cells^30,100^.

### Isodesmic polymerization within a condensate

How might the length of PopZ filaments be regulated, or the size of any polymer that forms isodesmically, in the context of a condensate? We hypothesize that the length distribution of filaments may be capped by the concentration in the dense phase. The thermodynamics of isodesmic polymerization implies that, as concentration increases, the population shifts to larger filament lengths. However, condensate thermodynamics dictates that the concentration in the dense phase (c_dense_) remains constant while its volume expands with increasing total protein concentration. Any condition that affects c_dense_ without interfering with filamentation may therefore affect the sampled filament lengths. This might explain the slight difference in filament lengths we observed in our cryo-ET data between WT and OD-only condensates (Fig. S1D, right panel). Similarly, clients may affect the packing of PopZ within condensates and shift the filament length distribution to smaller values by reducing c_dense_ (see next section).

### Material properties of the PopZ microdomain *in vivo*

We have demonstrated that the material properties of PopZ condensates are tightly coupled to its ability to form filaments. In previous work, we showed that PopZ variants altering the material properties of the PopZ microdomain in *Caulobacter crescentus* adversely affect fitness^30^. In the same work, we manipulated the sequence and length of the IDR to change material properties without interfering with filamentation. We propose that different species of α-proteobacteria may require PopZ microdomains with distinct material properties tailored to their unique physiology and environments. Evolutionarily, the simplest and most adaptable method to fine-tune these properties likely involves adjusting the IDR sequence rather than the highly conserved OD. This idea agrees with our observation that tweaking the IDR sequence reduced bacterial fitness (previous work^30^), while altering the ratio of hexamer-to-filaments by mutating the OD abolished growth altogether (this work, Figs. 6 and 7D). With respect to the mechanism by which changes in material properties harm *Caulobacter*, we hypothesize that viscosity is crucial for sequestration of clients within the microdomain, whereas surface tension may be important for compartmentalization and stability of the microdomain at the poles. The ability of PopZ to sequester clients and establish gradients of client proteins such as ParA and CtrA is crucial for asymmetric cell division^29^. Indeed, the client ChpT is sequestered less efficiently when the fraction of hexamers is increased, confirming that filamentation affects client retention (Fig. 3F). Additionally, high viscosity or increased client retention might stabilize protein:protein interactions^101^ and promote enzymatic reactions by reducing kinetics. Surface tension, on the other hand, increases the resistance to external forces and might help keep the microdomain in place and maintain its structural integrity under mechanical stress or changes in osmolarity. Higher surface tension creates a sharper boundary, which could make interactions more selective and spatially defined or generate forces that push or pull other condensates, such as the ParB:*parS* complex or DNA, during chromosome segregation.

Regarding the cross-linking of PopZ by other clients, we speculate that client proteins might change the condensate’s material properties depending on their expression levels. The length of isodesmic polymers within condensates is concentration-dependent, and cross-linking PopZ would alter the thermodynamics of the system, including c_dense_, therefore affecting filament length, which in turn affects material properties. If multiple clients compete for the same binding sites on PopZ, as is expected based on the literature^37^, but only some of them can form cross-links through dimerization or oligomerization, a scenario becomes plausible where cell-cycle dependent expression of clients correlates with changes in the material properties of the compartment.

### Future directions

Here, we provided a mechanistic model of PopZ assembly from monomers to trimers, hexamers, filaments, and ultimately, condensates. We described how H1-IDR prevents filaments from interacting and how overcoming this inhibition releases and extends H1-IDR, leading to interfilament contacts that drive condensates into which clients can partition. The material properties of these condensates can be affected drastically (by modifying the OD) or subtly (by modifying the IDR). Cryo-ET was instrumental in characterizing PopZ condensates, allowing us to bridge nanometer- to micrometer-scale insights, capturing both individual supramolecular structures and the broader condensate architecture.

Future studies will focus on developing new analytical methods tailored for cryo-ET to gain deeper insights into the structural and dynamic properties of condensates formed by interconnected filaments. In addition, the modular and tunable nature of PopZ, combined with our mechanistic model, provides a robust framework for rationally designing experiments or applications. Researchers can target specific stages along the assembly pathway to enhance or disrupt PopZ assembly, with potential applications across cell biology and synthetic biology.

## Acknowledgments

We thank members of the Lasker, Deniz, and Racki labs for insightful discussions. We thank Danielle Grotjahn and her lab members for their expert advice and assistance with cryo-ET data analysis. We also thank J Hammond, Phil Ordoukhanian, and the Biophysics and Biochemistry Core at Scripps Research for their support of biochemistry experiments. We gratefully acknowledge support from the NIH (NINDS DP2 NS142714 to S.B., NIGMS F32 GM150243 to A.P.L, NIGMS R01 GM083960 to A. Sali, NINDS R01 NS095892 and NIGMS RO1 GM14305 to G.C.L., NIGMS R35 GM130375 to A.A.D, and ORIPS10 OD032467 to Scripps Research for equipment). We are also gratefully acknowledging support from the NSF (2235200 to A. Salazar and DBI 2213983, Water and Life Interface Institute, to S.B. and A.S.H). K.L. acknowledges the Gordon & Betty Moore Foundation for support of this work through the Moore Inventor Fellowship number 579361. S.B. is a CPRIT scholar in Cancer Research and work in his lab is supported by CPRIT (RR220094).

## Author contributions

Conceptualization: D.S., A.A.D, K.L. Protein purification: D.S., A.K. Mass photometry, smFRET, AUC: D.S. Cryo-ET sample preparation T.B. Cryo-ET data processing and analysis: T.B., D.P. *In vitro* and *in vivo* imaging: D.S. Slab simulations: A.L., A. Sali. Cryo-EM sample preparation: A. Salazar. Cryo-EM data processing and analysis: A. Salazar, G.C.L. Formal analysis: D.S., S.B., A.S.H. K.L. Writing (original draft): D.S., K.L. Writing (review & editing): D.S., A.L., S.B., A.S.H., A. Sali, D.P., A.A.D, K.L.

## References

1. Banani, S.F., Lee, H.O., Hyman, A.A., and Rosen, M.K. (2017). Biomolecular condensates: organizers of cellular biochemistry. Nat Rev Mol Cell Biol 18, 285–298. 10.1038/nrm.2017.7.

2. Shin, Y., and Brangwynne, C.P. (2017). Liquid phase condensation in cell physiology and disease. Science 357. 10.1126/science.aaf4382.

3. Boeynaems, S., Chong, S., Gsponer, J., Holt, L., Milovanovic, D., Mitrea, D.M., Mueller-Cajar, O., Portz, B., Reilly, J.F., Reinkemeier, C.D., et al. (2023). Phase Separation in Biology and Disease; Current Perspectives and Open Questions. J Mol Biol 435, 167971. 10.1016/j.jmb.2023.167971.

4. Lyon, A.S., Peeples, W.B., and Rosen, M.K. (2021). A framework for understanding the functions of biomolecular condensates across scales. Nat Rev Mol Cell Biol 22, 215–235. 10.1038/s41580-020-00303-z.

5. Choi, J.M., Holehouse, A.S., and Pappu, R.V. (2020). Physical Principles Underlying the Complex Biology of Intracellular Phase Transitions. Annu Rev Biophys 49, 107–133. 10.1146/annurev-biophys-121219-081629.

6. Pei, G., Lyons, H., Li, P., and Sabari, B.R. (2024). Transcription regulation by biomolecular condensates. Nat Rev Mol Cell Biol. 10.1038/s41580-024-00789-x.

7. Su, X., Ditlev, J.A., Hui, E., Xing, W., Banjade, S., Okrut, J., King, D.S., Taunton, J., Rosen, M.K., and Vale, R.D. (2016). Phase separation of signaling molecules promotes T cell receptor signal transduction. Science 352, 595–599. 10.1126/science.aad9964.

8. Alberti, S., and Hyman, A.A. (2021). Biomolecular condensates at the nexus of cellular stress, protein aggregation disease and ageing. Nat Rev Mol Cell Biol 22, 196–213. 10.1038/s41580-020-00326-6.

9. Visser, B.S., Lipinski, W.P., and Spruijt, E. (2024). The role of biomolecular condensates in protein aggregation. Nat Rev Chem 8, 686–700. 10.1038/s41570-024-00635-w.

10. Pappu, R.V., Cohen, S.R., Dar, F., Farag, M., and Kar, M. (2023). Phase Transitions of Associative Biomacromolecules. Chem Rev 123, 8945–8987. 10.1021/acs.chemrev.2c00814.

11. Mittag, T., and Pappu, R.V. (2022). A conceptual framework for understanding phase separation and addressing open questions and challenges. Mol Cell 82, 2201–2214. 10.1016/j.molcel.2022.05.018.

12. Gouveia, B., Kim, Y., Shaevitz, J.W., Petry, S., Stone, H.A., and Brangwynne, C.P. (2022). Capillary forces generated by biomolecular condensates. Nature 609, 255–264. 10.1038/s41586-022-05138-6.

13. Alshareedah, I., Borcherds, W.M., Cohen, S.R., Singh, A., Posey, A.E., Farag, M., Bremer, A., Strout, G.W., Tomares, D.T., and Pappu, R.V. (2024). Sequence-specific interactions determine viscoelasticity and ageing dynamics of protein condensates. Nature Physics 20, 1482–1491.

14. Feric, M., Vaidya, N., Harmon, T.S., Mitrea, D.M., Zhu, L., Richardson, T.M., Kriwacki, R.W., Pappu, R.V., and Brangwynne, C.P. (2016). Coexisting Liquid Phases Underlie Nucleolar Subcompartments. Cell 165, 1686–1697. 10.1016/j.cell.2016.04.047.

15. Posey, A.E., Bremer, A., Erkamp, N.A., Pant, A., Knowles, T.P.J., Dai, Y., Mittag, T., and Pappu, R.V. (2024). Biomolecular Condensates are Characterized by Interphase Electric Potentials. J Am Chem Soc. 10.1021/jacs.4c08946.

16. Peran, I., and Mittag, T. (2020). Molecular structure in biomolecular condensates. Curr Opin Struct Biol 60, 17–26. 10.1016/j.sbi.2019.09.007.

17. Bienz, M. (2020). Head-to-Tail Polymerization in the Assembly of Biomolecular Condensates. Cell 182, 799–811. 10.1016/j.cell.2020.07.037.

18. Fletcher, D.A., and Mullins, R.D. (2010). Cell mechanics and the cytoskeleton. Nature 463, 485–492 10.1038/nature08908.

19. Berkamp, S., Mostafavi, S., and Sachse, C. (2021). Structure and function of p62/SQSTM1 in the emerging framework of phase separation. FEBS J 288, 6927–6941. 10.1111/febs.15672.

20. Bouchard, J.J., Otero, J.H., Scott, D.C., Szulc, E., Martin, E.W., Sabri, N., Granata, D., Marzahn, M.R., Lindorff-Larsen, K., Salvatella, X., et al. (2018). Cancer Mutations of the Tumor Suppressor SPOP Disrupt the Formation of Active, Phase-Separated Compartments. Mol Cell 72, 19–36 e18. 10.1016/j.molcel.2018.08.027.

21. Weirich, K.L., Banerjee, S., Dasbiswas, K., Witten, T.A., Vaikuntanathan, S., and Gardel, M.L. (2017). Liquid behavior of cross-linked actin bundles. Proc Natl Acad Sci U S A 114, 2131–2136. 10.1073/pnas.1616133114.

22. Bowman, G.R., Comolli, L.R., Zhu, J., Eckart, M., Koenig, M., Downing, K.H., Moerner, W.E., Earnest, T., and Shapiro, L. (2008). A polymeric protein anchors the chromosomal origin/ParB complex at a bacterial cell pole. Cell 134, 945–955. 10.1016/j.cell.2008.07.015.

23. Ebersbach, G., Briegel, A., Jensen, G.J., and Jacobs-Wagner, C. (2008). A self-associating protein critical for chromosome attachment, division, and polar organization in caulobacter. Cell 134, 956–968. 10.1016/j.cell.2008.07.016.

24. Bowman, G.R., Comolli, L.R., Gaietta, G.M., Fero, M., Hong, S.H., Jones, Y., Lee, J.H., Downing, K.H., Ellisman, M.H., McAdams, H.H., and Shapiro, L. (2010). Caulobacter PopZ forms a polar subdomain dictating sequential changes in pole composition and function. Mol Microbiol 76, 173–189. 10.1111/j.1365-2958.2010.07088.x.

25. Ettema, T.J., and Andersson, S.G. (2009). The alpha-proteobacteria: the Darwin finches of the bacterial world. Biol Lett 5, 429–432. 10.1098/rsbl.2008.0793.

26. Tsokos, C.G., and Laub, M.T. (2012). Polarity and cell fate asymmetry in Caulobacter crescentus. Curr Opin Microbiol 15, 744–750. 10.1016/j.mib.2012.10.011.

27. Govers, S.K., and Jacobs-Wagner, C. (2020). Caulobacter crescentus: model system extraordinaire. Curr Biol 30, R1151–R1158. 10.1016/j.cub.2020.07.033.

28. Lasker, K., Mann, T.H., and Shapiro, L. (2016). An intracellular compass spatially coordinates cell cycle modules in Caulobacter crescentus. Curr Opin Microbiol 33, 131–139. 10.1016/j.mib.2016.06.007.

29. Lasker, K., von Diezmann, L., Zhou, X., Ahrens, D.G., Mann, T.H., Moerner, W.E., and Shapiro, L. (2020). Selective sequestration of signalling proteins in a membraneless organelle reinforces the spatial regulation of asymmetry in Caulobacter crescentus. Nat Microbiol 5, 418–429. 10.1038/s41564-019-0647-7.

30. Lasker, K., Boeynaems, S., Lam, V., Scholl, D., Stainton, E., Briner, A., Jacquemyn, M., Daelemans, D., Deniz, A., Villa, E., et al. (2022). The material properties of a bacterial-derived biomolecular condensate tune biological function in natural and synthetic systems. Nature communications 13, 5643. 10.1038/s41467-022-33221-z.

31. Heebner, J.E., Purnell, C., Hylton, R.K., Marsh, M., Grillo, M.A., and Swulius, M.T. (2022). Deep Learning-Based Segmentation of Cryo-Electron Tomograms. J Vis Exp. 10.3791/64435.

32. Toro-Nahuelpan, M., Plitzko, J.M., Schüler, D., and Pfeiffer, D. (2022). In vivo Architecture of the Polar Organizing Protein Z (PopZ) Meshwork in the Alphaproteobacteria Magnetospirillum gryphiswaldense and Caulobacter crescentus. J Mol Biol 434, 167423. 10.1016/j.jmb.2021.167423.

33. Oosawa, F., and Kasai, M. (1962). A theory of linear and helical aggregations of macromolecules. J Mol Biol 4, 10–21. 10.1016/s0022-2836(62)80112-0.

34. Romberg, L., Simon, M., and Erickson, H.P. (2001). Polymerization of Ftsz, a bacterial homolog of tubulin. is assembly cooperative? J Biol Chem 276, 11743–11753. 10.1074/jbc.M009033200.

35. Bowman, G.R., Perez, A.M., Ptacin, J.L., Ighodaro, E., Folta-Stogniew, E., Comolli, L.R., and Shapiro, L. (2013). Oligomerization and higher-order assembly contribute to sub-cellular localization of a bacterial scaffold. Mol Microbiol 90, 776–795. 10.1111/mmi.12398.

36. Laloux, G., and Jacobs-Wagner, C. (2013). Spatiotemporal control of PopZ localization through cell cycle-coupled multimerization. J Cell Biol 201, 827–841. 10.1083/jcb.201303036.

37. Holmes, J.A., Follett, S.E., Wang, H., Meadows, C.P., Varga, K., and Bowman, G.R. (2016). Caulobacter PopZ forms an intrinsically disordered hub in organizing bacterial cell poles. Proc Natl Acad Sci U S A 113, 12490–12495. 10.1073/pnas.1602380113.

38. Ptacin, J.L., Gahlmann, A., Bowman, G.R., Perez, A.M., von Diezmann, A.R., Eckart, M.R., Moerner, W.E., and Shapiro, L. (2014). Bacterial scaffold directs pole-specific centromere segregation. Proc Natl Acad Sci U S A 111, E2046–2055. 10.1073/pnas.1405188111.

39. Soltermann, F., Foley, E.D.B., Pagnoni, V., Galpin, M., Benesch, J.L.P., Kukura, P., and Struwe, W.B. (2020). Quantifying Protein-Protein Interactions by Molecular Counting with Mass Photometry. Angew Chem Int Ed Engl 59, 10774–10779. 10.1002/anie.202001578.

40. Young, G., Hundt, N., Cole, D., Fineberg, A., Andrecka, J., Tyler, A., Olerinyova, A., Ansari, A., Marklund, E.G., Collier, M.P., et al. (2018). Quantitative mass imaging of single biological macromolecules. Science 360, 423–427. 10.1126/science.aar5839.

41. Howlett, G.J., Minton, A.P., and Rivas, G. (2006). Analytical ultracentrifugation for the study of protein association and assembly. Curr Opin Chem Biol 10, 430–436. 10.1016/j.cbpa.2006.08.017.

42. Agudo-Canalejo, J., Schultz, S.W., Chino, H., Migliano, S.M., Saito, C., Koyama-Honda, I., Stenmark, H., Brech, A., May, A.I., Mizushima, N., and Knorr, R.L. (2021). Wetting regulates autophagy of phase-separated compartments and the cytosol. Nature 591, 142–146. 10.1038/s41586-020-2992-3.

43. Alshareedah, I., Kaur, T., and Banerjee, P.R. (2021). Methods for characterizing the material properties of biomolecular condensates. Methods Enzymol 646, 143–183. 10.1016/bs.mie.2020.06.009.

44. Ditlev, J.A., Case, L.B., and Rosen, M.K. (2018). Who’s In and Who’s Out-Compositional Control of Biomolecular Condensates. J Mol Biol 430, 4666–4684. 10.1016/j.jmb.2018.08.003.

45. Nordyke, C.T., Ahmed, Y., Puterbaugh, R.Z., Bowman, G.R., and Varga, K. (2020). Intrinsically disordered bacterial polar organizing protein Z, PopZ, interacts with protein binding partners through an N-terminal Molecular Recognition Feature. J Mol Biol. 10.1016/j.jmb.2020.09.020.

46. Biondi, E.G., Reisinger, S.J., Skerker, J.M., Arif, M., Perchuk, B.S., Ryan, K.R., and Laub, M.T. (2006). Regulation of the bacterial cell cycle by an integrated genetic circuit. Nature 444, 899–904. 10.1038/nature05321.

47. Jumper, J., Evans, R., Pritzel, A., Green, T., Figurnov, M., Ronneberger, O., Tunyasuvunakool, K., Bates, R., Žídek, A., Potapenko, A., et al. (2021). Highly accurate protein structure prediction with AlphaFold. Nature 596, 583–589. 10.1038/s41586-021-03819-2.

48. Abramson, J., Adler, J., Dunger, J., Evans, R., Green, T., Pritzel, A., Ronneberger, O., Willmore, L., Ballard, A.J., Bambrick, J., et al. (2024). Accurate structure prediction of biomolecular interactions with AlphaFold 3. Nature 630, 493–500. 10.1038/s41586-024-07487-w.

49. Scholl, D., and Deniz, A.A. (2022). Conformational Freedom and Topological Confinement of Proteins in Biomolecular Condensates. J Mol Biol 434, 167348. 10.1016/j.jmb.2021.167348.

50. Saar, K.L., Qian, D., Good, L.L., Morgunov, A.S., Collepardo-Guevara, R., Best, R.B., and Knowles, T.P.J. (2023). Theoretical and Data-Driven Approaches for Biomolecular Condensates. Chem Rev 123, 8988–9009. 10.1021/acs.chemrev.2c00586.

51. Latham, A.P., and Zhang, B. (2022). Unifying coarse-grained force fields for folded and disordered proteins. Curr Opin Struct Biol 72, 63–70. 10.1016/j.sbi.2021.08.006.

52. Regy, R.M., Thompson, J., Kim, Y.C., and Mittal, J. (2021). Improved coarse-grained model for studying sequence dependent phase separation of disordered proteins. Protein Sci 30, 1371–1379. 10.1002/pro.4094.

53. Liu, S., Wang, C., Latham, A.P., Ding, X., and Zhang, B. (2023). OpenABC enables flexible, simplified, and efficient GPU accelerated simulations of biomolecular condensates. PLoS Comput Biol 19, e1011442. 10.1371/journal.pcbi.1011442.

54. Dignon, G.L., Zheng, W., Kim, Y.C., Best, R.B., and Mittal, J. (2018). Sequence determinants of protein phase behavior from a coarse-grained model. PLoS Comput Biol 14, e1005941. 10.1371/journal.pcbi.1005941.

55. Noel, J.K., Levi, M., Raghunathan, M., Lammert, H., Hayes, R.L., Onuchic, J.N., and Whitford, P.C. (2016). SMOG 2: A Versatile Software Package for Generating Structure-Based Models. PLoS Comput Biol 12, e1004794. 10.1371/journal.pcbi.1004794.

56. Bentley, E.P., Scholl, D., Wright, P.E., and Deniz, A.A. (2023). Coupling of binding and differential subdomain folding of the intrinsically disordered transcription factor CREB. FEBS Lett 597, 917–932. 10.1002/1873-3468.14554.

57. Lerner, E., Barth, A., Hendrix, J., Ambrose, B., Birkedal, V., Blanchard, S.C., Börner, R., Sung Chung, H., Cordes, T., Craggs, T.D., et al. (2021). FRET-based dynamic structural biology: Challenges, perspectives and an appeal for open-science practices. Elife 10. 10.7554/eLife.60416.

58. Fitzkee, N.C., and Rose, G.D. (2004). Reassessing random-coil statistics in unfolded proteins. Proc Natl Acad Sci U S A 101, 12497–12502. 10.1073/pnas.0404236101.

59. Lotthammer, J.M., Ginell, G.M., Griffith, D., Emenecker, R.J., and Holehouse, A.S. (2024). Direct prediction of intrinsically disordered protein conformational properties from sequence. Nat Methods 21, 465–476. 10.1038/s41592-023-02159-5.

60. Das, R.K., Ruff, K.M., and Pappu, R.V. (2015). Relating sequence encoded information to form and function of intrinsically disordered proteins. Curr Opin Struct Biol 32, 102–112. 10.1016/j.sbi.2015.03.008.

61. Antonik, M., Felekyan, S., Gaiduk, A., and Seidel, C.A. (2006). Separating structural heterogeneities from stochastic variations in fluorescence resonance energy transfer distributions via photon distribution analysis. J Phys Chem B 110, 6970–6978. 10.1021/jp057257+.

62. Livny, J., Yamaichi, Y., and Waldor, M.K. (2007). Distribution of centromere-like parS sites in bacteria: insights from comparative genomics. J Bacteriol 189, 8693–8703. 10.1128/JB.01239-07.

63. Ptacin, J.L., Lee, S.F., Garner, E.C., Toro, E., Eckart, M., Comolli, L.R., Moerner, W.E., and Shapiro, L. (2010). A spindle-like apparatus guides bacterial chromosome segregation. Nat Cell Biol 12, 791–798. 10.1038/ncb2083.

64. Shebelut, C.W., Guberman, J.M., van Teeffelen, S., Yakhnina, A.A., and Gitai, Z. (2010). Caulobacter chromosome segregation is an ordered multistep process. Proc Natl Acad Sci U S A 107, 14194–14198. 10.1073/pnas.1005274107.

65. Puentes-Rodriguez, S.G., Norcross, J.D., and Mera, P.E. (2023). To let go or not to let go: how ParA can impact the release of the chromosomal anchoring in Caulobacter crescentus. Nucleic Acids Res 51, 12275–12287. 10.1093/nar/gkad982.

66. Flory, P.J. (1953). Principles of polymer chemistry (Cornell university press).

67. Rubinstein, M., and Colby, R.H. (2003). Polymer physics (Oxford university press).

68. de Gennes, P. (1987). Polymers at an interface; a simplified view. Advances in colloid and interface science 27, 189–209.

69. Laghmach, R., Alshareedah, I., Pham, M., Raju, M., Banerjee, P.R., and Potoyan, D.A. (2022). RNA chain length and stoichiometry govern surface tension and stability of protein-RNA condensates. iScience 25, 104105. 10.1016/j.isci.2022.104105.

70. Pyo, A.G.T., Zhang, Y., and Wingreen, N.S. (2023). Proximity to criticality predicts surface properties of biomolecular condensates. Proc Natl Acad Sci U S A 120, e2220014120. 10.1073/pnas.2220014120.

71. Li, P., Banjade, S., Cheng, H.C., Kim, S., Chen, B., Guo, L., Llaguno, M., Hollingsworth, J.V., King, D.S., Banani, S.F., et al. (2012). Phase transitions in the assembly of multivalent signalling proteins. Nature 483, 336–340. 10.1038/nature10879.

72. Jain, A., and Vale, R.D. (2017). RNA phase transitions in repeat expansion disorders. Nature 546, 243–247. 10.1038/nature22386.

73. Langdon, E.M., Qiu, Y., Ghanbari Niaki, A., McLaughlin, G.A., Weidmann, C.A., Gerbich, T.M., Smith, J.A., Crutchley, J.M., Termini, C.M., Weeks, K.M., et al. (2018). mRNA structure determines specificity of a polyQ-driven phase separation. Science 360, 922–927. 10.1126/science.aar7432.

74. Lin, Y.H., Forman-Kay, J.D., and Chan, H.S. (2018). Theories for Sequence-Dependent Phase Behaviors of Biomolecular Condensates. Biochemistry 57, 2499–2508. 10.1021/acs.biochem.8b00058.

75. Vernon, R.M., Chong, P.A., Tsang, B., Kim, T.H., Bah, A., Farber, P., Lin, H., and Forman-Kay, J.D. (2018). Pi-Pi contacts are an overlooked protein feature relevant to phase separation. Elife 7. 10.7554/eLife.31486.

76. Wang, J., Choi, J.M., Holehouse, A.S., Lee, H.O., Zhang, X., Jahnel, M., Maharana, S., Lemaitre, R., Pozniakovsky, A., Drechsel, D., et al. (2018). A Molecular Grammar Governing the Driving Forces for Phase Separation of Prion-like RNA Binding Proteins. Cell 174, 688–699 e616. 10.1016/j.cell.2018.06.006.

77. Martin, E.W., Holehouse, A.S., Peran, I., Farag, M., Incicco, J.J., Bremer, A., Grace, C.R., Soranno, A., Pappu, R.V., and Mittag, T. (2020). Valence and patterning of aromatic residues determine the phase behavior of prion-like domains. Science 367, 694–699. 10.1126/science.aaw8653.

78. Wu, H., and Fuxreiter, M. (2016). The Structure and Dynamics of Higher-Order Assemblies: Amyloids, Signalosomes, and Granules. Cell 165, 1055–1066. 10.1016/j.cell.2016.05.004.

79. Goetz, S.K., and Mahamid, J. (2020). Visualizing Molecular Architectures of Cellular Condensates: Hints of Complex Coacervation Scenarios. Dev Cell 55, 97–107. 10.1016/j.devcel.2020.09.003.

80. Bienz, M. (2014). Signalosome assembly by domains undergoing dynamic head-to-tail polymerization. Trends Biochem Sci 39, 487–495. 10.1016/j.tibs.2014.08.006.

81. Ciuffa, R., Lamark, T., Tarafder, A.K., Guesdon, A., Rybina, S., Hagen, W.J., Johansen, T., and Sachse, C. (2015). The selective autophagy receptor p62 forms a flexible filamentous helical scaffold. Cell Rep 11, 748–758. 10.1016/j.celrep.2015.03.062.

82. Jakobi, A.J., Huber, S.T., Mortensen, S.A., Schultz, S.W., Palara, A., Kuhm, T., Shrestha, B.K., Lamark, T., Hagen, W.J.H., Wilmanns, M., et al. (2020). Structural basis of p62/SQSTM1 helical filaments and their role in cellular cargo uptake. Nature communications 11, 440. 10.1038/s41467-020-14343-8.

83. Zaffagnini, G., Savova, A., Danieli, A., Romanov, J., Tremel, S., Ebner, M., Peterbauer, T., Sztacho, M., Trapannone, R., Tarafder, A.K., et al. (2018). p62 filaments capture and present ubiquitinated cargos for autophagy. Embo J 37. 10.15252/embj.201798308.

84. Sun, D., Wu, R., Zheng, J., Li, P., and Yu, L. (2018). Polyubiquitin chain-induced p62 phase separation drives autophagic cargo segregation. Cell Res 28, 405–415. 10.1038/s41422-018-0017-7.

85. Goode, B.L., and Eck, M.J. (2007). Mechanism and function of formins in the control of actin assembly. Annu Rev Biochem 76, 593–627. 10.1146/annurev.biochem.75.103004.142647.

86. Wang, K., and Singer, S.J. (1977). Interaction of filamin with f-actin in solution. Proc Natl Acad Sci U S A 74, 2021–2025. 10.1073/pnas.74.5.2021.

87. Yin, H.L., and Stossel, T.P. (1979). Control of cytoplasmic actin gel-sol transformation by gelsolin, a calcium-dependent regulatory protein. Nature 281, 583–586. 10.1038/281583a0.

88. Yin, H.L., Hartwig, J.H., Maruyama, K., and Stossel, T.P. (1981). Ca2+ control of actin filament length. Effects of macrophage gelsolin on actin polymerization. J Biol Chem 256, 9693–9697.

89. Kadzik, R.S., Homa, K.E., and Kovar, D.R. (2020). F-Actin Cytoskeleton Network Self-Organization Through Competition and Cooperation. Annu Rev Cell Dev Biol 36, 35–60. 10.1146/annurev-cellbio-032320-094706.

90. Cuneo, M.J., and Mittag, T. (2019). The ubiquitin ligase adaptor SPOP in cancer. FEBS J 286, 3946–3958. 10.1111/febs.15056.

91. Clark, A., and Burleson, M. (2020). SPOP and cancer: a systematic review. Am J Cancer Res 10, 704–726.

92. Wang, Z., Song, Y., Ye, M., Dai, X., Zhu, X., and Wei, W. (2020). The diverse roles of SPOP in prostate cancer and kidney cancer. Nat Rev Urol 17, 339–350. 10.1038/s41585-020-0314-z.

93. Marzahn, M.R., Marada, S., Lee, J., Nourse, A., Kenrick, S., Zhao, H., Ben-Nissan, G., Kolaitis, R.M., Peters, J.L., Pounds, S., et al. (2016). Higher-order oligomerization promotes localization of SPOP to liquid nuclear speckles. Embo J 35, 1254–1275. 10.15252/embj.201593169.

94. Cuneo, M.J., O’Flynn, B.G., Lo, Y.H., Sabri, N., and Mittag, T. (2023). Higher-order SPOP assembly reveals a basis for cancer mutant dysregulation. Mol Cell 83, 731–745 e734. 10.1016/j.molcel.2022.12.033.

95. Thomasen, F.E., Cuneo, M.J., Mittag, T., and Lindorff-Larsen, K. (2023). Conformational and oligomeric states of SPOP from small-angle X-ray scattering and molecular dynamics simulations. Elife 12. 10.7554/eLife.84147.

96. Zhang, Q., Shi, Q., Chen, Y., Yue, T., Li, S., Wang, B., and Jiang, J. (2009). Multiple Ser/Thr-rich degrons mediate the degradation of Ci/Gli by the Cul3-HIB/SPOP E3 ubiquitin ligase. Proc Natl Acad Sci U S A 106, 21191–21196. 10.1073/pnas.0912008106.

97. Pierce, W.K., Grace, C.R., Lee, J., Nourse, A., Marzahn, M.R., Watson, E.R., High, A.A., Peng, J., Schulman, B.A., and Mittag, T. (2016). Multiple Weak Linear Motifs Enhance Recruitment and Processivity in SPOP-Mediated Substrate Ubiquitination. J Mol Biol 428, 1256–1271. 10.1016/j.jmb.2015.10.002.

98. Kwon, J.E., La, M., Oh, K.H., Oh, Y.M., Kim, G.R., Seol, J.H., Baek, S.H., Chiba, T., Tanaka, K., Bang, O.S., et al. (2006). BTB domain-containing speckle-type POZ protein (SPOP) serves as an adaptor of Daxx for ubiquitination by Cul3-based ubiquitin ligase. J Biol Chem 281, 12664–12672. 10.1074/jbc.M600204200.

99. Schmit, J.D., Bouchard, J.J., Martin, E.W., and Mittag, T. (2020). Protein Network Structure Enables Switching between Liquid and Gel States. J Am Chem Soc 142, 874–883. 10.1021/jacs.9b10066.

100. Boeynaems, S., Dorone, Y., Zhuang, Y., Shabardina, V., Huang, G., Marian, A., Kim, G., Sanyal, A., Şen, N.-E., and Griffith, D. (2023). Poly (A)-binding protein is an ataxin-2 chaperone that regulates biomolecular condensates. Mol Cell 83, 2020–2034. e2026.

101. Elbaum-Garfinkle, S., Kim, Y., Szczepaniak, K., Chen, C.C., Eckmann, C.R., Myong, S., and Brangwynne, C.P. (2015). The disordered P granule protein LAF-1 drives phase separation into droplets with tunable viscosity and dynamics. Proc Natl Acad Sci U S A 112, 7189–7194. 10.1073/pnas.1504822112.

