## supplementary figures and methods for "Cellular Function of a Biomolecular Condensate Is Determined by Its Ultrastructure"

Affiliations:
[1] Department of Integrative Structural and Computational Biology, The Scripps Research Institute, La Jolla, CA, 92037, USA
[2] Quantitative Biosciences Institute, University of California, San Francisco, San Francisco, CA 94158, USA.
[3] Department of Bioengineering and Therapeutic Sciences, University of California, San Francisco, San Francisco, CA 94158, USA.
[4] Department of Pharmaceutical Chemistry, University of California, San Francisco, San Francisco, CA 94158, USA
[5] Center for Infectious Disease Imaging, National Institutes of Health, Clinical Center, 10 Center Drive, Bethesda, MD 20892, USA
[6] Department of Molecular and Human Genetics, Baylor College of Medicine, Houston, TX 77030, USA
[7] Jan and Dan Duncan Neurological Research Institute, Texas Children's Hospital, Houston, TX 77030, USA
[8] Therapeutic Innovation Center (THINC), Baylor College of Medicine, Houston, TX 77030, USA
[9] Center for Alzheimer's and Neurodegenerative Diseases (CAND), Texas Children's Hospital, Houston, TX 77030, USA
[10] Dan L Duncan Comprehensive Cancer Center (DLDCCC), Baylor College of Medicine, Houston, TX 77030, USA
[11] Department of Biochemistry and Molecular Biophysics, Washington University School of Medicine, St. Louis, MO
[12] Center for Biomolecular Condensates (CBC), Washington University in St. Louis, St. Louis, MO

#### Supplementary Figures

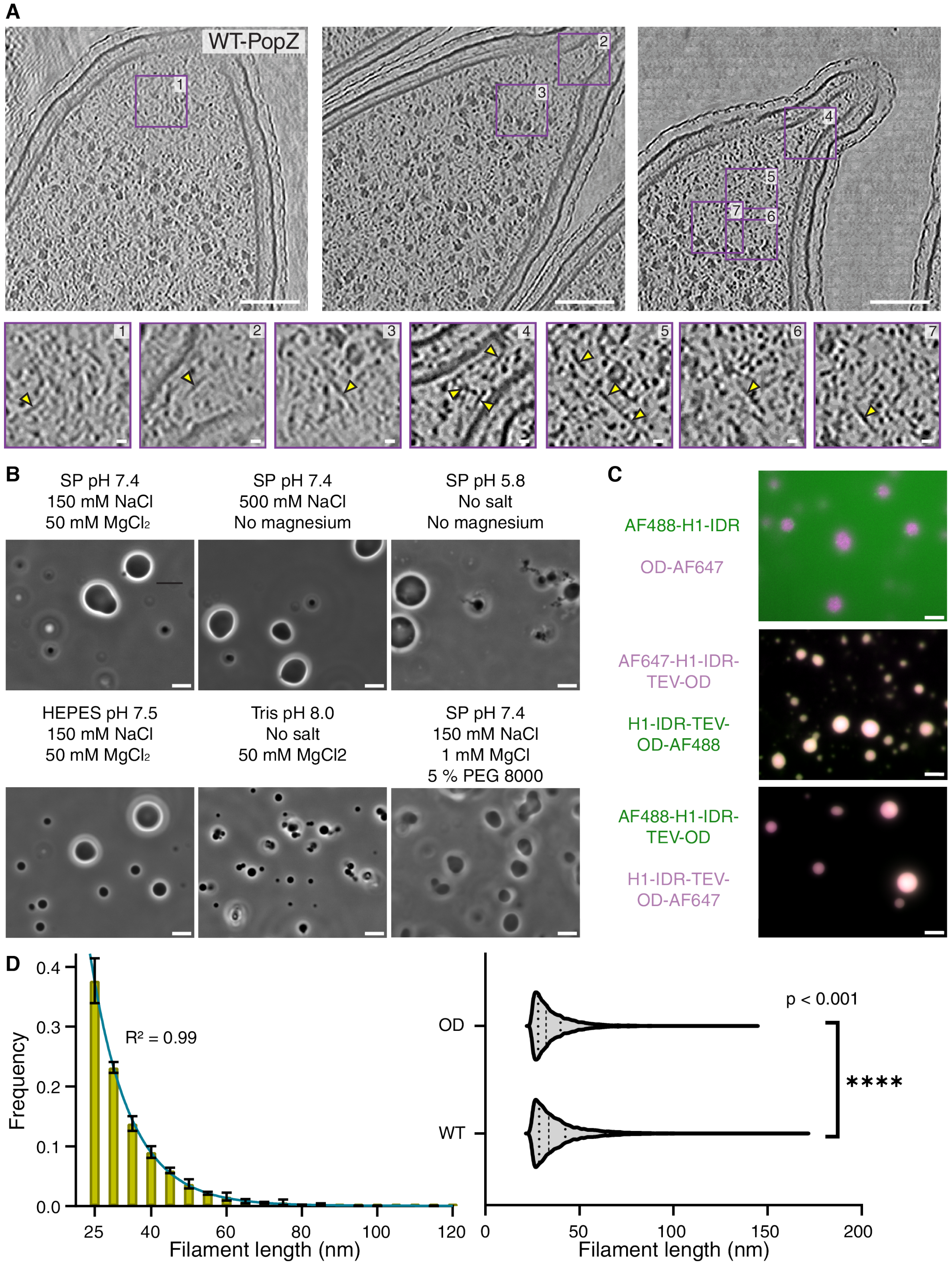

**Figure S1. Condensation and filamentation of full-length PopZ and its oligomerization domain**

**(A)** Cryo-ET of *Caulobacter* cells reveals filaments within PopZ microdomains. Shown are three representative microdomains in ∆*popZ Caulobacter* cells expressing mCherry-tagged

WT-PopZ. Scale bars: 100 nm. The purple squares indicate zoomed-in regions. Yellow arrows highlight filamentous structures. Scale bars: 10 nm. **(B)** PopZ condensation requires $MgCl_2$, high salt, or low *p*H. Phase contrast images of 5 $\mu$M purified PopZ in the annotated buffer conditions. Scale bar: 5 $\mu$m. **(C)** Fluorescence microscopy of PopZ constructs labeled with AF647 and AF488 attached either to Cys-20 (N-term) or Cys132 (OD). (upper two panels) Both uncleaved constructs readily partition into WT-PopZ droplets formed using 50 mM $MgCl_2$, regardless of the dye positions. (lower panel) The isolated OD forms droplets, whereas the H1-IDR remains in solution, regardless of the dye position. Scale bars: 5 $\mu$m. **(D)** Filament length distribution comparison between isolated OD and FL PopZ. (left) Distribution of the curved length of OD filaments. Columns indicate the mean filament length across three condensates; error bars denote standard deviation. (right) Violin plot comparison between the curved length distributions of the isolated OD and the FL protein. Dotted lines represent the median and quartiles. A two-tailed Mann-Whitney U test was used to evaluate statistical significance.

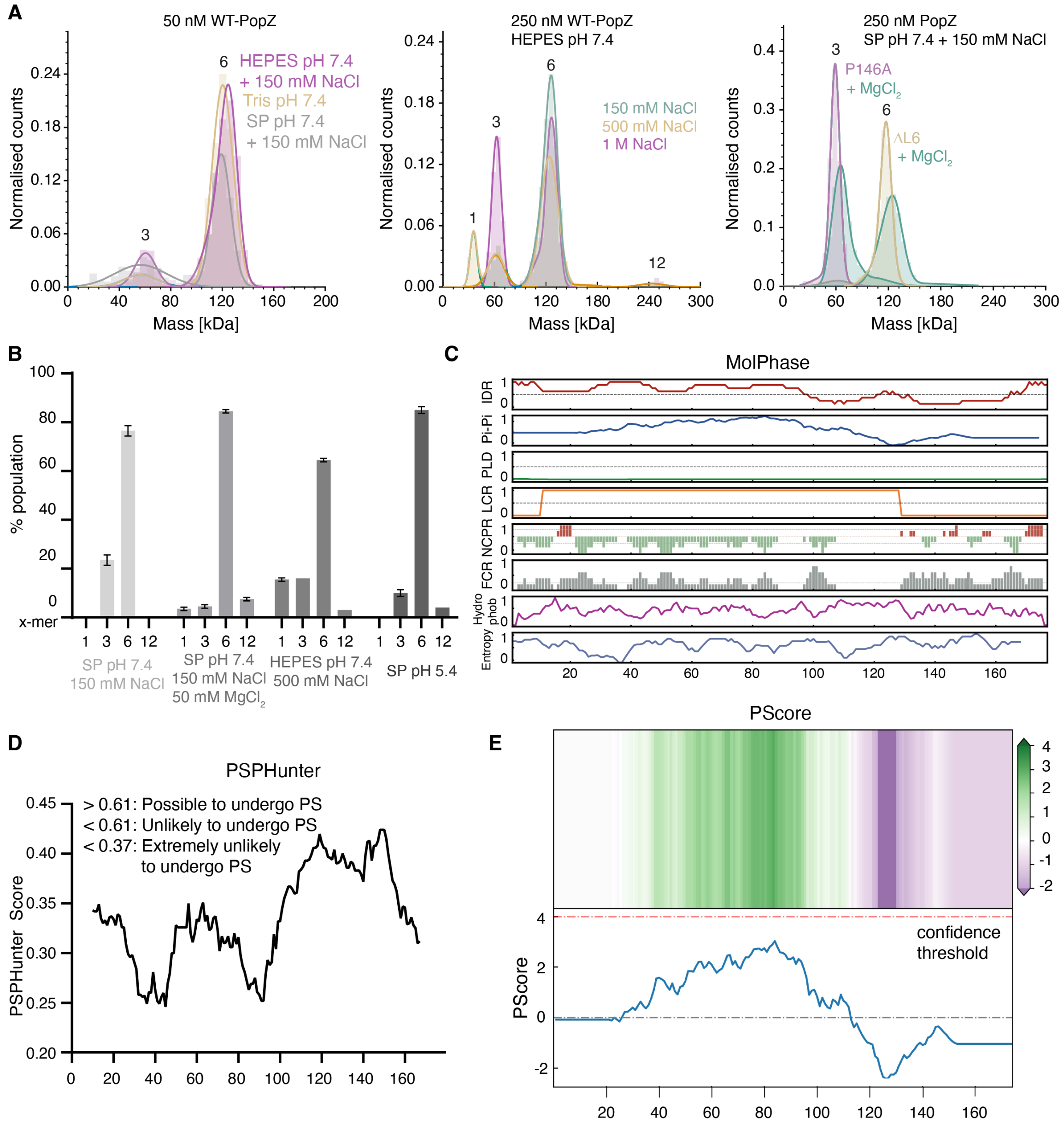

**Figure S2. Oligomerization of PopZ in various conditions**

**(A)** Mass photometry analysis of WT-PopZ and variants in different buffers. (left) 50 nM WT-PopZ assemble into trimers and hexamers in different buffers. (middle) Effect of salt concentration on PopZ oligomerization. (right) Addition of 50 mM MgCl2 did not affect the oligomerization of P146A and ΔL6. **(B)** Relative populations of oligomers measured by mass photometry using 250 nM PopZ. Conditions that trigger condensation at higher concentrations lead to the stacking of hexamers. Columns represent the mean of biological duplicates; error bars represent the standard deviation. **(C-E)** Graphs of per-residue analyses of the full-length PopZ sequence and scores from three of the mentioned phase separation predictors.

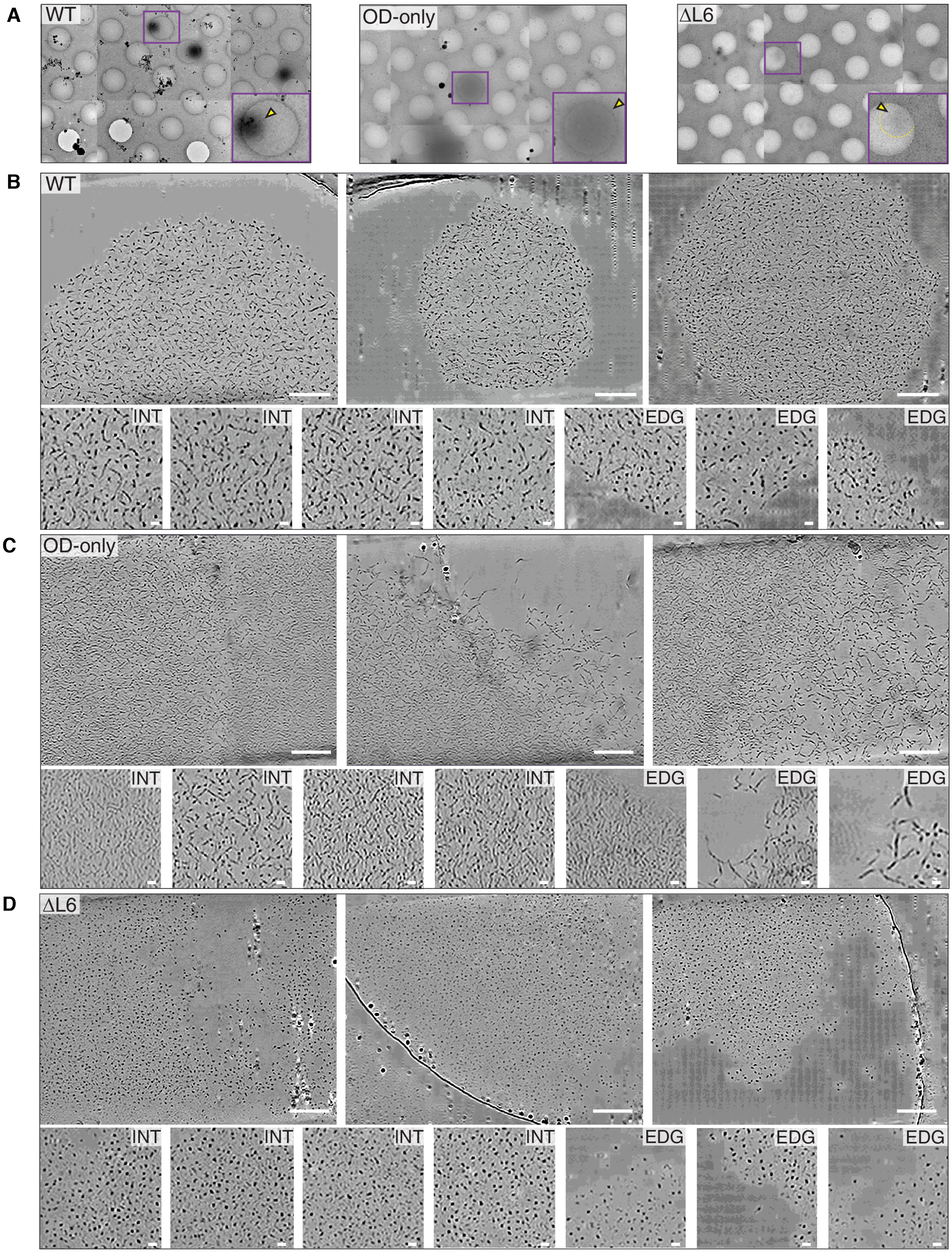

**Figure S3.** ***In vitro*** **condensates of WT, OD-only, and ΔL6-PopZ**
**(A)** Representative cryo-ET images at 3600X magnification of WT, OD-only, and DL6 PopZ condensates. The purple boxes indicate zoomed-in regions. The yellow arrow points to the condensate within the image. For DL6, the boundary of the condensate is blurry and indicated by the yellow dotted line. **(B-D)** Upper panels: Three representative cryo-ET images of WT, OD-only, and DL6 PopZ condensates (Scale bar: 100 nm). Lower panels, zoomed-in images of the condensates. Scale bar: 10 nm. 'INT' indicates that the image was taken from the interior of the condensate, and 'EDG' indicates that the image was taken from the edge.

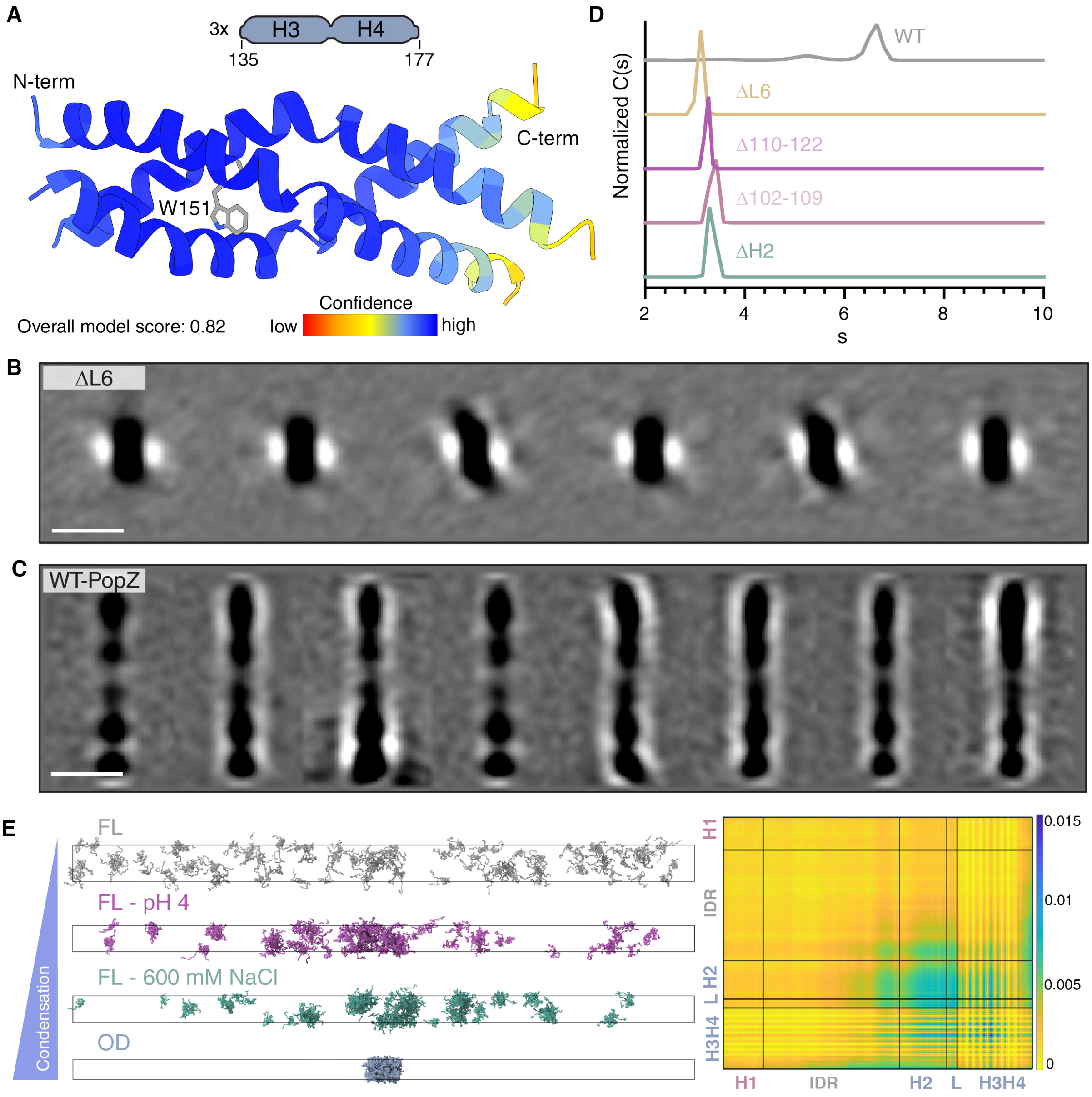

**Figure S4. Sequence determinants for oligomerization, filamentation, and condensation**
**(A)** AlphaFold 2 prediction for H3H4 trimer. The top-ranking model, with an overall score of 0.82, is colored by pLDDT. **(B)** 2D cross-sections of six class averages for ΔL6-PopZ. Scale bar: 10 nm. **(C)** 2D cross-sections of class average structures for WT-PopZ. Scale bar: 10 nm. **(D)** Deletion of H2 or its fragments traps PopZ as a hexamer. Analytical ultracentrifugation of H2 deletion variants at approximately 100 $\mu$M in 10 mM sodium phosphate *p*H 7.4 + 150 mM NaCl. **(E)** Simulation results agree with our experimental observation and point to interaction

regions between PopZ trimers. (left) The last frames of slab simulations for the indicated conditions. (right) Inter-trimer contact map of FL-PopZ at 600 mM NaCl with color coding indicating contact frequency. Residue by residue contact maps were calculated with a cutoff distance of 1 nm in MDAnalysis[1,2].

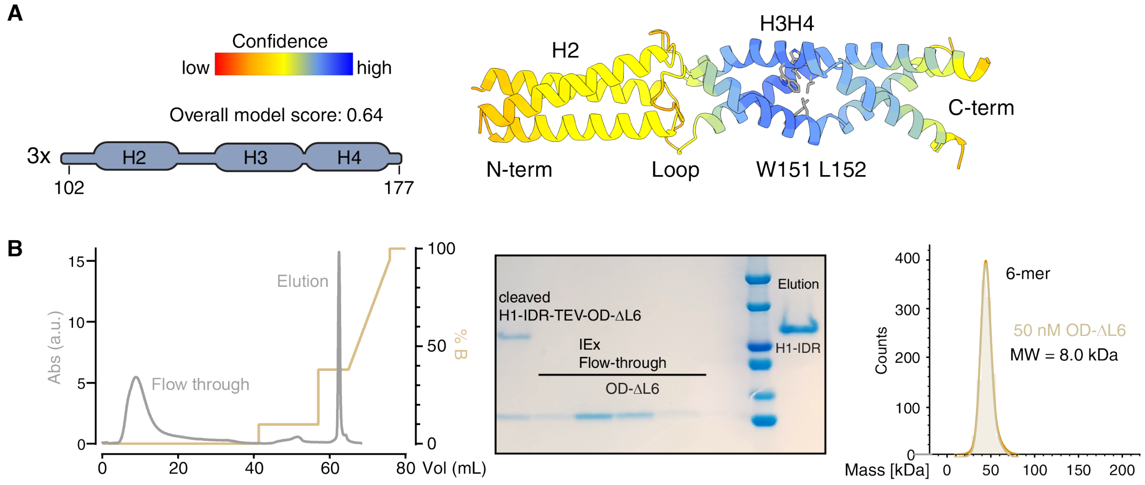

**Figure S5. Trimer model and purification of isolated ΔL6-OD**

**(A)** AlphaFold2 prediction for ΔL6-OD trimer. The top-ranking model of H2-loop-H3H4 trimer, with an overall score of 0.64, is colored by pLLDT. **(B)** (left) Ion-exchange chromatogram of H1-IDR-TEV-OD-ΔL6 after TEV cleavage. (middle) SDS-PAGE gel showing that OD-ΔL6 stayed in the ion-exchange flow-through, whereas H1-IDR attached to the column and was concentrated in the elution fraction. (right) Mass photometry confirms that isolated OD-ΔL6 forms hexamers.

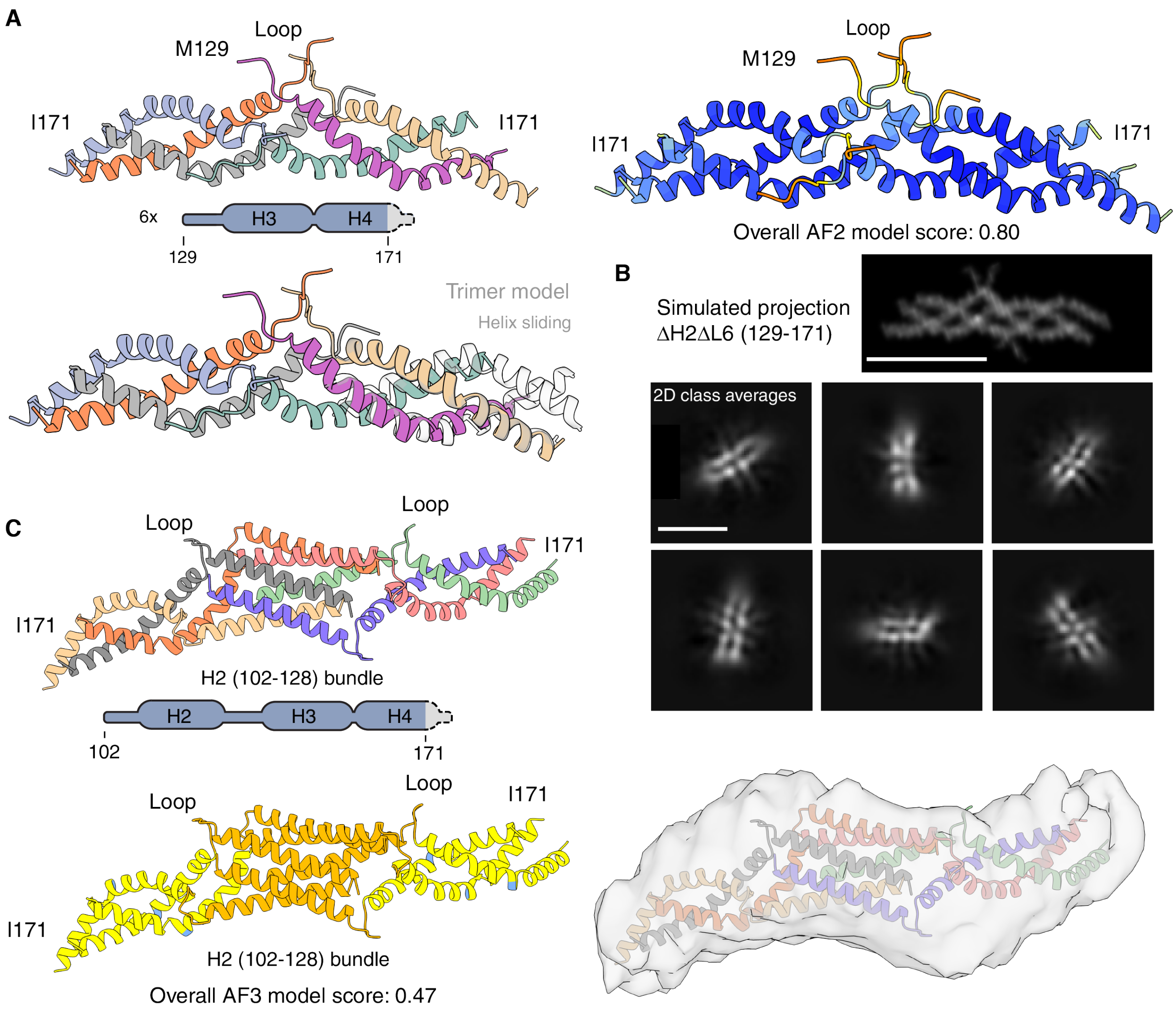

**Figure S6. Structural models of PopZ hexamers**

**(A)** (top) AlphaFold 2 hexamer model using six 'loop-H3H4ΔL6' chains, colored by chain (left) and pLDDT score (right). (bottom) Overlay of the same hexamer with the model of the trimer in transparent gray. **(B)** (top) Simulated projection of the hexamer model of the ΔH2ΔL6-PopZ. AlphaFold prediction was converted to a molecular envelope using the ChimeraX molmap command, and the resulting density was shown using volumetric rendering. (bottom) Cryo-EM 2D class averages of ΔH2ΔL6-PopZ show a similar elongated organization. Scale bars: 60 Å. **(C)** AlphaFold 3 model of six H2-loop-H3H4ΔL6 chains, colored by chain (top) and pLDDT score (bottom). The structural model fits into the low-resolution envelope of the curved hexamer 2D class obtained via cryo-ET sub-tomogram averaging (bottom right).

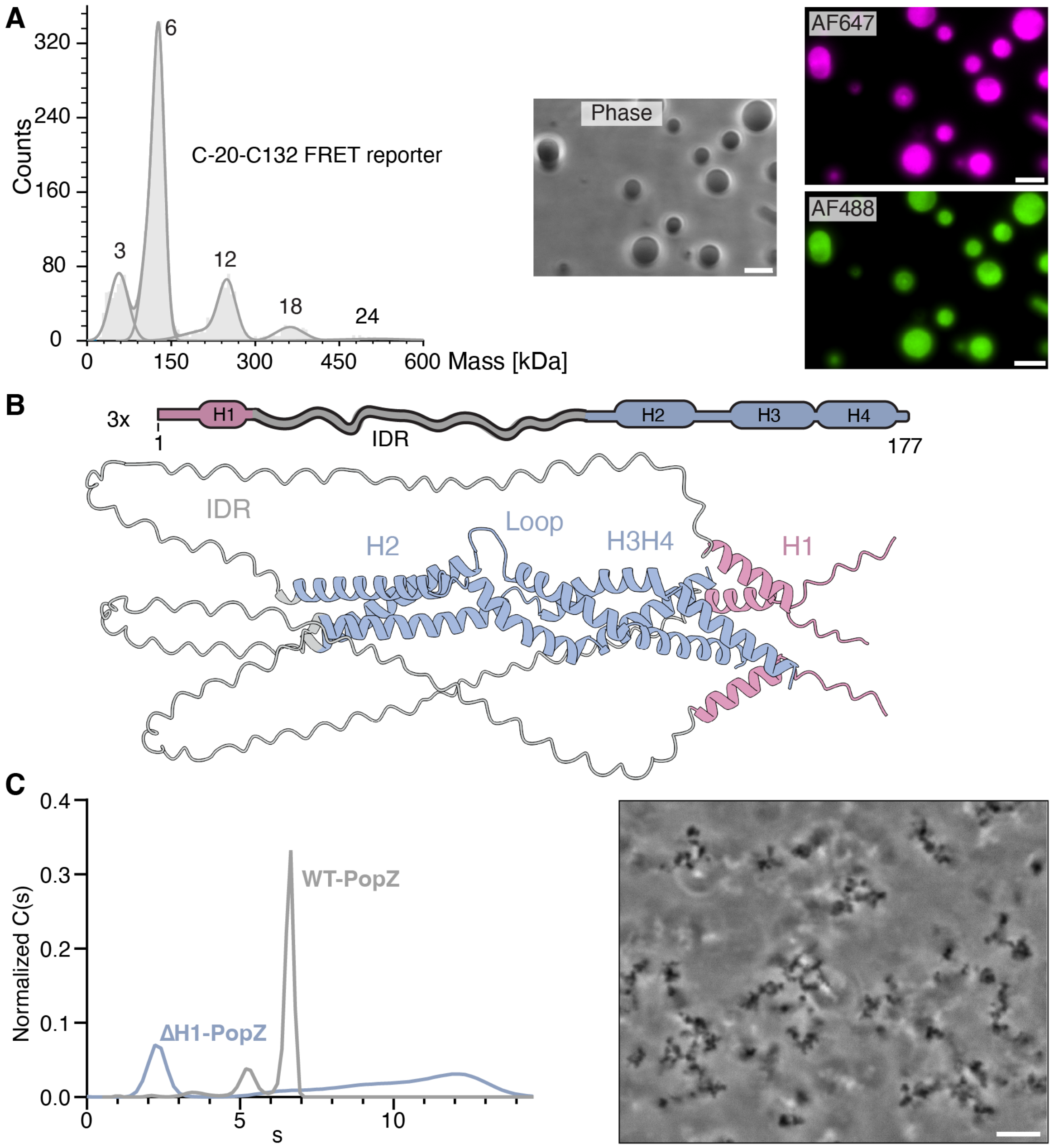

**Figure S7. The H1 region interacts with the OD**
**(A)** Labeled PopZ behaves like WT-PopZ. (left) Mass photometry of the smFRET reporter labeled at C-20 and C132 (illustrated in Fig. 2A) shows that the labeled reporter adopts the same oligomeric species as WT-PopZ upon MgCl2 addition. (right) Fluorescence microscopy shows that the labeled FRET reporter forms spherical droplets, akin to WT-PopZ. **(B)** AF3 model of three FL-PopZ molecules. **(C)** Characterization of ΔH1-PopZ. (left) AUC shows that in the absence of MgCl2, ΔH1-PopZ forms larger species than the WT filaments. (right) Phase imaging of 70 $\mu$M ΔH1-PopZ in 10 mM sodium phosphate pH 7.4, 150 mM NaCl, 50 mM MgCl2 shows formation of aggregate-like clusters.

### Tables

#### Table S1: Phase separation predictors

| PopZ sequence | FuzDrop[3] | MolPhase[4] | PScore[5] | PSPHunter[6] | PicNic[7] | PSPredictor[8] |
|---|---|---|---|---|---|---|
| **FL** | Y(0.980) | Y(0.995) | N(1.63) | N(0.377) | N(0.315) | Y(0.662) |
| **H1-IDR** | Y(0.977) | Y(0.948) | / | N(0.372) | / | Y(0.944) |
| **OD** | N(0.100) | N(0.004) | / | N(0.047) | / | N(0.006) |

While some of the phase separation predictors correctly predicted that full-length (FL) PopZ condenses, they falsely attributed this to the H1-IDR. No predictor captured the ability of the oligomerization domain (OD) to condense. 'Y' means the phase separation predictor predicts condensation, 'N' means it predicts no condensation, and '/' means it could not be applied because the sequence was below the minimum length requirement.

#### Table S2: Oligomerization, filamentation, and condensation behavior of PopZ variants

| Variant | Csat (μM) | Alternative Condition*1 | Oligomerization in Mass Photometry | Filamentation in AUC |
|---|---|---|---|---|
| **WT** | < 1 | / | 1,3,6,12,18,24 | Yes |
| **ΔL6** | ~ 20*2 | *p*H 6.0, no salt, 50 mM $MgCl_2$ | 1, 3, 6 | No |
| **P146A** | ~ 2 | / | 1, 3 | Yes |
| **Δ129-134 (=Δloop)** | ~ 7 | / | 1, 3 | / |
| **Δ133-177** | / | / | 1 | / |
| **Δ102-128 (=ΔH2)** | / | 1 M NaCl 200 mM $MgCl_2$ | 1, 3, 6 | No |
| **Δ102-109** | / | 1 M NaCl 200 mM $MgCl_2$ | 1, 3, 6 | No |
| **Δ110-122** | / | 1 M NaCl 200 mM $MgCl_2$ | 1, 3, 6 | No |
| **P146A-W151G-L152S** | / | / | 1 | / |
| **Isolated OD-ΔL6** | ~10 | | | |

*1 Some variants did not form droplets under our reference condition of 10 mM sodium phosphate *p*H 7.4 + 150 mM NaCl and 50 mM $MgCl_2$. The 'Alternative Condition' column specifies a different condition under which we could observe droplets, if applicable.

*[2] ΔL6 droplets formed under our reference condition were too small for FRAP experiments. We used an alternative buffer (10 mM sodium phosphate *p*H 6.0, 50 mM $MgCl_2$) for those experiments. In this buffer, the $c_{sat}$ of ΔL6 is below 10 µM.

#### Table S3: Fluorescence recovery after photobleaching

| Simple exponential | AF647-PopZ | | | AF488-ChpT | | |
|---|---|---|---|---|---|---|
| | Recovery $\tau_{1/2}$ | Mobile fraction | $R^2$ | Recovery $\tau_{1/2}$ | Mobile fraction | $R^2$ |
| **5 *µ*M WT-PopZ** | 1076 ± 910 s | 5.5 ± 2.2 % | 0.24 | 208 ± 17 s | 67.1 ± 1.6 % | 0.98 |
| **2.5 *µ*M WT-PopZ + 5 *µ*M ΔL6-PopZ** | 1866 ± 685 s | 44.5 ± 10.2 % | 0.94 | 137 ± 16 s | 75.6 ± 2.0 % | 0.97 |
| **10 *µ*M ΔL6-PoPZ** | 596 ± 17 s | 90.8 ± 1.0 % | 0.99 | 97 ± 13 s | 88.8 ± 2.2 ± | 0.97 |

| Two-phase | AF488-ChpT | | | | |
|---|---|---|---|---|---|
| | Recovery $\tau_{1/2}$ | | | Mobile fraction | $R^2$ |
| | % fast | Fast | Slow | | |
| **5 *µ*M WT-PopZ** | 34 ± 3 % | 34 ± 6 s | 351 ± 61 s | 76.3 ± 2.6 % | 0.99 |
| **2.5 *µ*M WT-PopZ + 5 *µ*M ΔL6-PopZ** | 37 ± 6 % | 19 ± 7 s | 216 ± 46 s | 81.9 ± 2.3 % | 0.99 |
| **10 *µ*M ΔL6-PoPZ** | 40 ± 5 % | 13 ± 4 s | 159 ± 29 s | 95.0 ± 2.1 % | 0.99 |

The shown values correspond to a 95% confidence interval.

### Methods

#### Media and growth conditions for bacterial in vivo studies

*Caulobacter* strains were grown at 30 °C in PYE or M2G liquid media supplemented with kanamycin (5 *µ*g/mL). When indicated, cells were synchronized as follows[9]: 30 mL of bacteria in log phase ($OD_{600}$ ~0.5) were pelleted, resuspended in 1 mL cold M2, repelleted and resuspended in 900 *µ*L M2 and 900 *µ*L Percoll (Cytiva). They were then pelleted for 20 min at 15,000 g and 4 °C to separate swarmer and stalked cells. Swarmer cells were transferred into M2 and washed three times. Finally, the bacteria were resuspended in PYE and diluted to an OD of 0.12 and transferred to a pad of PYE and 1.2% agarose. Expression of PopZ variants was indirectly induced through the leaky expression from PYE media[10].

#### PopZ expression and purification

PopZ was expressed and purified from *E. coli* strain BL21. Cells were grown in 0.5 L LB kanamycin (50 *µ*g/mL) to an OD of 0.4 at 37 °C and then switched to induction temperatures of 18 °C for 20 min prior to induction. PopZ expression was induced overnight with 0.1 mM IPTG. Cell pellets were collected via centrifugation and stored at –80 °C.

The entire purification was done under denaturing conditions of 8 M urea to prevent aggregation and condensation of PopZ. The frozen cell pellet was resuspended in approximately 100 mL lysis buffer containing 10 mM sodium phosphate pH 7.2, 10 mM Tris-HCl pH 8.0, 300 mM NaCl, 8 M urea, 20 mM imidazole, and 1 tablet of EDTA-free protease inhibitors (Pierce) for every 50 mL of lysis buffer. Cells were lysed by sonication and insoluble material was removed via centrifugation at 12,000 g for 30 min at 12 °C. The supernatant was injected onto a 5 mL HisTrap FF (Cytiva) via an AKTA Go system (Cytiva). The supernatant was reinjected once. PopZ was eluted through a step gradient of imidazole. The eluted PopZ was diluted 4-fold in ion-exchange buffer A, containing 10 mM sodium phosphate pH 7.2, 10 mM Tris-HCl pH 8.0, 50 mM NaCl, 8 M urea. The diluted sample was then loaded onto a 1 mL HiTrap Q FF (Cytiva) ion-exchange column and eluted with a salt gradient. The purified protein was flash frozen and stored at -80°C. It was buffer exchanged into the desired buffer immediately before experiments. Purity of PopZ variants was assessed via SDS-PAGE and mass spectrometry.

#### Cryo-electron tomography

##### Grid Preparation for in vitro samples

50-200 $\mu$L of 10 nm gold fiducials (Aurion) were spun down in a benchtop centrifuge for 20 min at 15,000 rpm. The supernatant was removed and the beads were washed twice with standard buffer (10 mM sodium phosphate pH 7.4, 150 mM NaCl, 50 mM $MgCl_2$). They were then resuspended in an appropriate amount of buffer resulting in a 20% higher bead concentration than the original gold fiducial concentration. Quantifoil R 2/1 copper 200 mesh grids were glow-discharged with a Pelco easiGlow (Ted Pella, 25 s glow at 15 mA). 2.3 $\mu$L of gold fiducial beads were deposited onto the grid. The beads were given 30 seconds to spread out, then 4 $\mu$L of the PopZ sample was deposited onto the grid, back-side blotted, and plunged into a mixture of propane/ethane using either a Vitrobot Mark IV (Thermo Fisher Scientific; 2.5 s blot time, 0 s wait time, 0.5 s drain time, 0 blot force) or a homemade gravity-driven plunging device. After freezing, the vitrified grids were clipped with autogrids (Thermo Fisher Scientific).

##### Grid Preparation for *Caulobacter crescentus* samples

*ΔpopZ C. crescentus* strain expressing mCherry-WT-PopZ via a pXMCS-2 plasmid was grown at 30 °C in PYE Kanamycin until an $OD_{600}$ between 0.2 and 0.5, then diluted 100-fold and grown until an $OD_{600}$ of 0.3. Cells were then pelleted and resuspended in PYE to an $OD_{600}$ of 3. 5 $\mu$L of the sample was deposited on glow-discharged (Pelco easiGlow; 25 seconds glow at 15mA) Quantifoil R 2/1 copper 200 mesh grids. The grids were then back-blotted for 7-12 seconds before plunge freezing with a Vitrobot Mark IV into a mixture of 37% propane/ethane. After freezing, the vitrified grids were clipped with autogrids.

##### Focus-Ion beam milling

*C. crescentus* samples were subjected to cryo-FIB milling to reduce sample thickness and increase image contrast. Cryo-FIB milling was performed using the Aquilos 2 dualbeam cryo-FIB/SEM instrument (Thermo Fisher Scientific). The sample was sputtered with metallic platinum for 15 seconds, followed by a layer of organometallic platinum for 40 seconds, and then sputtered again with metallic platinum for 15 seconds. The metallic platinum helps to reduce charging and drifting while milling, and the organometallic platinum helps to protect the sample from the focused ion beam. Targeted regions on the grid were milled to produce lamellae with a thickness of approximately 150-200 nm. Finally, the grid was sputtered with metallic platinum for 15-20 seconds. The final sputter coat adds bead-like fiducial inclusions that are used for tilt-series alignment.

**Data collection**

Both *in vitro* PopZ samples and cryo-lamellae of *Caulobacter* samples were imaged with a Titan Krios 300 keV microscope (Thermo Fisher Scientific) equipped with a field emission gun, a direct detection device (Gatan K3), and an energy filter (operated with a slit width of 20 eV). The SerialEM[11] package with PACEtomo[12] scripts was used to collect image stacks at tilt angles ranging from +51° to -51° (3° increment) using a dose-symmetric scheme with a cumulative dose of ~105 e–/Å². Images were taken at a magnification resulting in 1.663 Å/pixel and a nominal defocus of ~-5 $\mu$m. Image stacks containing ~10 frames were motion-corrected with MotionCor2[13], and assembled and aligned by IMOD[14]. All tomograms used in the figures were denoised with CryoCARE[15] followed by Isonet[16].

**Subtomogram averaging**

For *in vitro* PopZ data, 4x binned tomograms were imported into EMAN2[17]. Straight segments of PopZ filaments were selected using a segmentation-based particle picking function. Particle positions were translated into the IMOD coordinate system to be imported into the i3[18] subtomogram averaging software. Before averaging, the CTF of tilted images was estimated using gCTF[19], CTF correction was performed using the IMOD ctfphaseflip function, and CTF-corrected tomograms were created using IMOD. An initial reference was created by manually picking ~100 straight segments of filaments with pre-defined orientations using Tomopick[20] software and averaging them using i3. Using this initial reference, the remaining particles were aligned and classified.

**Segmentation and filament tracing**

Denoised *in vitro* PopZ tomograms were segmented using the Deep Learning tool in Dragonfly[21] (Comet Technologies Canada Inc). 2D UNet models were trained on hand-segmented frames of the tomogram The model was applied to the entire tomogram, and significant false positives were manually removed from the 3D renderings. The island removal feature in Dragonfly was used to eliminate independent densities smaller than the size of a PopZ hexamer, thereby reducing noise from the segmentation results.

*In vitro* PopZ segmentations were used in Amira (Thermo Fisher Scientific) for filament tracing. The parameters for cylinder correlation were derived from measurements of PopZ filaments in IMOD and are as follows: cylinder length, 100; angular sampling, 5; mask cylinder radius, 30; outer cylinder radius, 27; inner cylinder radius, 0. Filaments shorter than 250 Å were excluded because filaments shorter than 250 Å tend to be fragmented parts of longer filaments. The parameters used for the trace correlation lines are as follows: minimum seed correlation, 50; minimum continuation, 100; direction coefficient, 0.4; minimum distance, 75; minimum length, 250; search cone length, 60; search cone angle, 45; minimum step size, 5. Tracing results were exported into .xml files containing the filament tracing metadata table. Parameters such as curved lengths and chord lengths were imported into Prism for further analysis. Statistical significance was determined using the Mann-Whitney U test, and the histogram of curved lengths was analyzed using a nonlinear regression (curve fit) approach.

**Data availability**
Representative tomograms generated in this study are available in the Electron Microscopy Data Resource under the following accession codes: EMD-47539 for an *in vitro* WT-PopZ condensate, EMD-47540 for an *in vitro* OD-PopZ condensate, EMD-47542 for an *in vitro* ΔL6-PopZ condensate, and EMD-47557 for a PopZ condensate at the pole of a *Caulobacter* cell. Unprocessed tilt series are available in the Electron Microscopy Public Image Archive (EMPIAR) under accession codes EMPIAR-12372 for a PopZ microdomain in Δ*popZ Caulobacter* cells expressing mCherry-tagged WT-PopZ, and EMPIAR-12373 of an *in vitro* PopZ condensate.

#### Single-particle cryo-electron microscopy

**Sample preparation**
Samples were kept at 4°C throughout the entire preparation process. ΔH2ΔL6-PopZ was prepared at 2 mg/ml in a buffer containing 10 mM sodium phosphate pH 7.4 and 150 mM NaCl. Quantifoil 300 mesh R0.6/1 UltrAuFoil Holey Gold Films were glow discharged under vacuum for 30 seconds at 15 mA in a Pelco easiGlow 91000 Glow Discharge Cleaning System (Ted Pella, Inc.). Three microliters of sample were applied to the surface of the grid and blot-plunged using a manual plunger in a 4 °C cold room with ~95% humidity. Grids were manually blotted with Whatman 1 filter paper for 10 seconds and immediately plunged into a liquid ethane pool cooled by liquid nitrogen. Grids were then clipped into AutoGrids (Thermo Fisher Scientific) under liquid nitrogen vapor and stored in liquid nitrogen until the day of imaging.

**Cryo-EM data acquisition**
Cryo-EM data were collected on a Thermo-Fisher Talos Arctica transmission electron microscope operating at 200 keV, which had been aligned for parallel illumination settings with a 30 μm condenser2 aperture. Micrographs were collected using a Falcon 4i electron detector, with an applied total electron exposure of 50 e-/Å$^2$. The EPU data collection software was used to collect micrographs at 190,000x nominal magnification (0.74 Å/pixel at the specimen level) with a nominal defocus range set to 0.8 μm - 1.4 μm under focus. A maximum image shift of eight μm with aberration-free image shift (AFIS) was used to obtain a single exposure in the center of the 0.6 μm holes.

**Image processing for ΔH2-PopZ**
A total of 8,762 micrographs were collected and processed in real-time using cryoSPARC Live (V4.5.3). Patch Motion Correction was used to mitigate motions and radiation damage that occur during imaging. Motion-corrected images were processed collectively at the end of data collection, beginning with CTF estimation using

CTFFIND4. The 7,042 micrographs with reported CTF resolutions better than 8 Å were selected for further processing, and 5,899,512 particle selections were identified using a blob picker with default parameters and a minimum particle diameter of 70 Å and a maximum particle diameter of 100 Å. Particle selections that had local power between 426.0 & 967.0 were used for subsequent processing, leaving a total of 3,595,338 particles that were extracted using a box size of 194-pixels. 2D classification of this extracted stack into 75 classes was performed using default parameters with the exception of 40 online EM iterations and 400 batch size per class.

#### Light microscopy

Epifluorescence microscopy was performed on a Nikon Ti2-E automated inverted microscope with a 100x oil immersion objective (CFI60 plan apochromat lens, N.A. 1.45; Nikon) and a Kinetix sCMOS camera (Teledyne Photometrics). Images were acquired with the NIS Elements software version 5.42.03 and further processed in ImageJ or MicrobeJ. Bacteria were grown in liquid media to an OD 0.1-0.3 and put on pads (1.2% agarose in PYE) secured between coverglass and No. 1.5 coverslips via silicone gaskets (VWR). mCherry was excited with the 594 nm epi line, at 12% intensity and imaged with 100 ms exposure time using a 603-800 nm emission filter. CFP was excited with the 440 nm line at 15% intensity and imaged with 800 ms exposure time using a 464-486 nm emission filter.

For FRAP experiments, the same instrument was used but images were acquired using a TIRF iLAS2 module (Gataca Systems). Before the experiments, unlabeled WT- and ΔL6-PopZ were mixed with either labeled ChpT or labeled PopZ and incubated for 12 hours before condensation was triggered to allow the sample to equilibrate. After droplets formed, they were allowed to settle until the intensity of the whole image was stable, usually 2 hours. Droplets were partially bleached in their center using a point stimulation. Data was corrected for photobleaching and time-dependent changes in intensity by including unbleached reference droplets and then normalized. The data was fit in GraphPad Prism to a simple exponential for labeled PopZ and a two-phase model for labeled ChpT.

#### TEV cleavage experiments and SDS-PAGE

In vitro TEV cleavage was initiated by adding 100 nM TEV protease (purified in-house) to unlabeled WT-PopZ at 5 $\mu$M in 50 mM Tris-HCl, pH 8.0 with 0.5 mM EDTA, 1 mM DTT. When fluorescently-labeled PopZ was also present, it was added at low nanomolar concentrations. Cleavage was allowed to proceed for 10 minutes unless otherwise specified.

For gel analysis, droplets were pelleted (2,000g for 10 min) and separated from the supernatant. Both fractions were run on a 4-20% Mini-PROTEAN TGX™ gradientgel (BioRad) and protein was revealed via a Coomassie blue type LC6060 SimplyBlue™ stain (Invitrogen).

#### Mass photometry

Mass photometry experiments were performed on a Refeyn TwoMP (Refeyn Ltd, Oxford, UK) calibrated with a mix of BSA (Sigma-Aldrich) and thyroglobulin (Sigma-

Aldrich, St. Louis, MO). Coverslips (WillCo Wells, Amsterdam, NL) and gaskets (Grace Bio Labs, Bend, OR, USA) were prepared by washing with dd$H_2O$ followed by isopropanol and again with dd$H_2O$, repeated three times, and then dried. 20 µL of buffer was added to each well to focus the instrument, then 15 µL of buffer was removed and replaced with 15 µL of sample and mixed by pipette for 2 s before frame acquisition. Frames were acquired over 60 s using AcquireMP (version 2022 R1, Refeyn Ltd, Oxford, UK) using standard settings. Data were processed and analysed by fitting a Gaussian distribution to the data using DiscoverMP (version 2022 R1, Refeyn Ltd, Oxford, UK).

#### Analytical ultracentrifugation

Sedimentation velocity measurements were done with the indicated protein concentrations (between 30 and 140 $\mu$M) in a solution of 10 mM sodium phosphate pH 7.4 + 150 mM NaCl at 25°C on a Beckman Optima XL-I instrument equipped with a AN-60 Ti rotor and at 40,000 RPM. Samples were monitored at 280 nm for 150 scans with 8 min intervals for a total of 20 h. Scans were processed using SEDFIT software[22]. Fitting parameters such as the buffer density (1.00555 g/mL), buffer viscosity (0.010016 poise), and partial specific volume (varied across PopZ constructs) were calculated by SEDNTERP[23]. Sedimentation velocity profiles are shown at a confidence level of 95%.

#### AlphaFold predictions

To generate structural models, we ran either AlphaFold-Multimer[24] v2.3.2 (AF2, from https://github.com/google-deepmind/alphafold) or AlphaFold3[25] (AF3, from alphafoldserver.com) on the following sequences:

| Domains | Residues | Sequence | AlphaFold version |
|---|---|---|---|
| **3x H3H4** | 3x 135–177 | TLEDVVRELLRPLLKEWLDQNLPRI VETKVEEEVQRISRGRGA | 2 |
| **3x H2-loop-H3H4** | 3x 102–128 | EVAEQLVGVSAASAAASAFGSLSS ALLMPKDGRTLEDVVRELLRPLLKE WLDQNLPRIVETKVEEEVQRISRGR GA | 2 |
| **6x loop-H3H4ΔL6** | 6x 129–171 | MPKDGRTLEDVVRELLRPLLKEWL DQNLPRIVETKVEEEVQRI | 2 |
| **6x H2-loop-H3H4ΔL6** | 6x 102–171 | EVAEQLVGVSAASAAASAFGSLSS ALL MPKDGRTLEDVVRELLRPLLKEWL DQNLPRIVETKVEEEVQRI | 3 |
| **3x FL-PopZ** | 3x 1–177 | MSDQSQEPTMEEILASIRRIISEDDA PAEPAAEAAPPPPPEPEPEPVSFD DEVLELTDPIAPEPELPPLETVGDID VYSPPEPESEPAYTPPPAAPVFDR DEVAEQLVGVSAASAAASAFGSLS SALLMPKDGRTLEDVVRELLRPLLK | 3 |

| | |
|---|---|
| | EWLDQNLPRIVETKVEEEVQRISRG RGA |

#### Simulations

##### Force field

For each PopZ sequence of interest, we created an initial atomic structural model of a trimer using AlphaFold-Multimer[24]. This initial model served as a template for computing structure-based potentials[26], which restrained the backbone and tertiary structure (default parameter values from the maximum entropy optimized force field (MOFF)[27] were used). Tertiary interactions were considered only within each H1 helix (residues 10-24 in WT PopZ), within each H2 helix (residues 102-128 in WT PopZ) or to stabilize the trimer formed by H3 and H4 (residues 136-171 in WT PopZ). Inter-molecular interactions were scored by the hydrophobicity scale model parameterized with the Urry hydrophobicity scale (HPS-Urry)[28]. All simulations were conducted using Debye Huckle electrostatics with 150 mM salt, unless otherwise noted. In the low pH (pH4) simulation, changes in pH were approximated by modulating amino acid residue charges. Specifically, histidine (old charge: 0.25, new charge: 1.0), aspartic acid (old charge: -1, new charge: 0.0), and glutamic acid (old charge: -1.0, new charge: 0.0) parameters were changed, while other residues were unchanged. Simulations were performed in OpenMM (version 7.5.1)[29], using the OpenABC software package (version 1.06)[30]. Examples are available at https://github.com/alatham13/Open_ABC_PopZ.

##### Simulation procedure and analysis

Slab simulations were performed for each PopZ variant and solution condition[31]. For each simulation, 100 PopZ trimers were initially placed in a 100 × 100 × 100 nm$^3$ simulation box. We then performed steepest descent energy minimization, followed by a 0.1 $\mu$s constant-temperature, constant-pressure simulation at 150 K and 1 bar using a Langevin integrator with a time coupling constant of 1 ps and a Monte Carlo barostat. This simulation resulted in a single dense phase. Then, the z-dimension of the simulation box was expanded by approximately 20 times the original size to 500 nm, resulting in a droplet with a dilute phase on either side. With this modified simulation box, we performed a constant-temperature, constant-volume simulation for 0.1 $\mu$s with a time coupling constant of 100 ps. During the simulation, we raised the temperature from 150 K to 300 K in steps of 1.5 K every 1 ns. The resulting equilibrated system was simulated for an additional 5 $\mu$s at constant-temperature of 300 K and constant-volume. The analysis relied only on the last 3 $\mu$s of the trajectory, sampled every 1 ns.

#### PopZ Labeling for Fluorescence Experiments

Single- or double cysteine constructs of PopZ were labeled with Alexa Fluor 488 and Alexa Fluor 647 maleimide dyes (Thermo Fisher Scientific, Waltham, MA, USA). Before labeling, IMAC-pure PopZ at 60–90 $\mu$M was buffer exchanged into labeling buffer (50 mM HEPES pH 7.2, 1 mM TCEP). To avoid preferential labelling by one dye over the other, substoichiometric additions of the dye mixture were made to a purified protein construct over 3 hours to a final six-fold molar excess of each dye. Then the labeling was allowed to proceed overnight. To quench the labeling reaction and exchange the sample back into denaturing buffer prior to ion-exchange clean-up, we buffer exchanged it into10 mM sodium phosphate pH 7.2, 10 mM Tris-HCl pH 8.0, 50 mM NaCl, 8 M urea, 2 mM DTT. It was then purified via ion-exchange, flash frozen and stored at -80 °C. To assess the purity, successful labeling, and integrity of labeled PopZ, it was subjected to mass photometry and light microscopy, where it behaved like unlabeled PopZ (Fig. S6A).

#### Single-molecule Förster Resonance Energy Transfer

##### Data recording

For each measurement, PopZ labeled at positions C-20 and C132 with AF488 and AF647 was diluted to 300 pM and supplemented with 5 $\mu$M unlabeled PopZ. Background or scatter samples were prepared similarly but without labeled protein. For each experiment, 1–4 h datasets were recorded at 22 °C on a homebuilt multiparameter fluorescence detection microscope with pulsed interleaved excitation (MFD-PIE). Emission from a pulsed 483-nm laser diode (LDH-D-C-485, PicoQuant, Berlin, Germany) was cleaned up (Semrock, Rochester, New York, FF01-482/25-25), emission from a 635-nm laser diode (LDH-D-C-640, PicoQuant, Berlin, Germany) was cleaned up (Semrock, Rochester, New York, FF01-635/ 18-25), and both lasers were alternated at 30 MHz using a waveform generator (Keysight), a picosecond delayer (Micro Photon Devices, Bolzano, Italy) connected to the laser drivers (PDL 800-D). The red laser was delayed by ~ 20 ns with respect to the blue laser. Linear polarization was cleaned up (Glan-Taylor Polarizer, Thorlabs, Newton, NJ, USA, GT10A) and the red and blue light were combined into a single mode optical fiber (kineFlex, Excelitas, Waltham, MA, USA) before the light (90 µW of 483 nm light and 70 µW of 635 nm light) was reflected into the back port of the microscope (Axiovert 200, Zeiss, Oberkochen, Germany) and to the objective (C-APOCHROMAT, 40×/1.2W, Zeiss, Oberkochen, Germany). Sample emission was transmitted through a polychroic mirror (Chroma, Bellows Falls, VT, USA, ZT488/640rpc), focused through a 75 µm pinhole and spectrally split (Semrock, FF593-Di03-25×36). The blue range was filtered (Semrock, Rochester, New York, FF03- 525/50-25), and polarization was split (PBS101, Thorlabs) into parallel and perpendicular channels. The red range was also filtered (Semrock, Rochester, New York, FF01-698/70- 25), and polarization was split (PBS101, Thorlabs). Photons were detected on four avalanche photodiodes (SPCM-AQR- 14, PerkinElmer, Waltham, MA, for the green parallel and perpendicular channels and for the red parallel channel and SPCM-AQRH-14, Excelitas, Waltham, MA, USA, for the red perpendicular channel), which were connected to a time-correlated single-photon counting (TCSPC) device (Multi- Harp 150 N, PicoQuant, Berlin, Germany). Signals were stored in 32-bit first-in-first-out (FIFO) files. Microscope alignment was verified using fluorescence correlation spectroscopy (FCS) on freely diffusing Alexa Fluor 488 and Alexa Fluor 647 (Thermo Fisher Scientific, Waltham, MA, USA). Instrument response functions (IRFs) were recorded one detector at-a-time in a solution of ATTO 488-CA or ATTO 655-CA in near-saturated centrifuged potassium iodide at a 25-kHz average count rate for a total of 25 × $10^6$ photons. Macrotime-dependent microtime shifting was corrected for two (blue/parallel and red/perpendicular) of four avalanche photodiodes (APDs) using the IRF data as input.

##### Data analysis

Data were analyzed with the PAM software[32] via standard procedures for MFD-PIE smFRET burst analysis[33,34]. Signals from each TCSPC routing channel (corresponding to the individual detectors) were divided in time gates to discern 483-nm excited FRET photons from 635-nm excited acceptor photons. A two-color MFD all-photon burst search algorithm using a 500-$\mu$s sliding time window (minimum of 100 photons per burst, minimum of 5 photons per time window) was used to identify single donor- and/or acceptor-labeled molecules in the fluorescence traces. Double labelled single molecules were selected from the raw burst data using a kernel density estimator (ALEX-2CDE < 12)[35], that also excluded other artefacts. Sparse slow-diffusing aggregates were removed from the data by excluding bursts exhibiting a burst duration > 12 ms except for conditions with condensates. Data was corrected in this order to obtain the absolute stoichiometry parameter S and absolute FRET efficiency E: background subtraction, donor emission crosstalk correction, acceptor direct excitation correction and relative detection efficiency correction. The last value depends on the used dye combination, but also on the used detectors. By making histograms of E versus measurement time, we corroborated that the distribution of E was invariant over the duration of the measurement.

**Photon distribution analysis**

Static PDA was carried out to obtain the absolute interdye distance distribution assuming two Gaussian distributed states[36]. For each FRET data set, raw bursts were re-binned in 1 ms time bins, and histograms were constructed and analyzed. Only bins with at least 20 and maximally 200 photons (to reduce calculation time) were used for PDA analysis. A two-state model for a Gaussian distance distribution was used to generate a library of simulated $E_{\mathrm{FRET}}$ values, which was fitted to the experimental $E_{\mathrm{FRET}}$ histogram using a reduced $\chi^2$–guided simplex search algorithm to obtain the amplitude, mean distance $R$ and width $\sigma$ of all Gaussian distributed substates and the area fraction of each state, $A$ (%). A probability density function (PDF) was calculated per state using the R and $\sigma$ parameters that describe the underlying Gaussian distributed states. The summed PDF was scaled to a total area of unity, with the PDF area of each state scaled to the corresponding fraction of molecules. A Jacobian was used to estimate confidence intervals (95%) of the fit parameters. Criteria for a good fit were a low reduced $\chi 2$ value as well as a weighted-residual plot free of trends.

### Supplementary References

1. Michaud-Agrawal, N., Denning, E.J., Woolf, T.B., and Beckstein, O. (2011). MDAnalysis: a toolkit for the analysis of molecular dynamics simulations. J Comput Chem *32*, 2319-2327. 10.1002/jcc.21787.
2. Gowers, R.J., Linke, M., Barnoud, J., Reddy, T.J.E., Melo, M.N., Seyler, S.L., Domanski, J., Dotson, D.L., Buchoux, S., and Kenney, I.M. (2019). MDAnalysis: a Python package for the rapid analysis of molecular dynamics simulations. Los Alamos National Laboratory (LANL), Los Alamos, NM (United States). 2575-9752.
3. Hatos, A., Tosatto, S.C.E., Vendruscolo, M., and Fuxreiter, M. (2022). FuzDrop on AlphaFold: visualizing the sequence-dependent propensity of liquid-liquid phase separation and aggregation of proteins. Nucleic Acids Res *50*, W337-W344. 10.1093/nar/gkac386.

4. Liang, Q., Peng, N., Xie, Y., Kumar, N., Gao, W., and Miao, Y. (2024). MolPhase, an advanced prediction algorithm for protein phase separation. Embo J *43*, 1898-1918. 10.1038/s44318-024-00090-9.
5. Vernon, R.M., Chong, P.A., Tsang, B., Kim, T.H., Bah, A., Farber, P., Lin, H., and Forman-Kay, J.D. (2018). Pi-Pi contacts are an overlooked protein feature relevant to phase separation. Elife *7*. 10.7554/eLife.31486.
6. Sun, J., Qu, J., Zhao, C., Zhang, X., Liu, X., Wang, J., Wei, C., Wang, M., Zeng, P., Tang, X., et al. (2024). Precise prediction of phase-separation key residues by machine learning. Nature communications *15*, 2662. 10.1038/s41467-024-46901-9.
7. Hadarovich, A., Singh, H.R., Ghosh, S., Rostam, N., Hyman, A.A., and Toth-Petroczy, A. (2023). PICNIC identifies condensate-forming proteins across organisms. bioRxiv, 2023.2006. 2001.543229.
8. Chu, X., Sun, T., Li, Q., Xu, Y., Zhang, Z., Lai, L., and Pei, J. (2022). Prediction of liquid-liquid phase separating proteins using machine learning. BMC Bioinformatics *23*, 72. 10.1186/s12859-022-04599-w.
9. Tsai, J.W., and Alley, M.R. (2001). Proteolysis of the Caulobacter McpA chemoreceptor is cell cycle regulated by a ClpX-dependent pathway. J Bacteriol *183*, 5001-5007.
10. Bowman, G.R., Perez, A.M., Ptacin, J.L., Ighodaro, E., Folta-Stogniew, E., Comolli, L.R., and Shapiro, L. (2013). Oligomerization and higher-order assembly contribute to sub-cellular localization of a bacterial scaffold. Mol Microbiol *90*, 776-795. 10.1111/mmi.12398.
11. Mastronarde, D.N. (2005). Automated electron microscope tomography using robust prediction of specimen movements. J Struct Biol *152*, 36-51. 10.1016/j.jsb.2005.07.007.
12. Eisenstein, F., Yanagisawa, H., Kashihara, H., Kikkawa, M., Tsukita, S., and Danev, R. (2023). Parallel cryo electron tomography on in situ lamellae. Nat Methods *20*, 131-138. 10.1038/s41592-022-01690-1.
13. Zheng, S.Q., Palovcak, E., Armache, J.P., Verba, K.A., Cheng, Y., and Agard, D.A. (2017). MotionCor2: anisotropic correction of beam-induced motion for improved cryo-electron microscopy. Nat Methods *14*, 331-332. 10.1038/nmeth.4193.
14. Mastronarde, D.N., and Held, S.R. (2017). Automated tilt series alignment and tomographic reconstruction in IMOD. J Struct Biol *197*, 102-113. 10.1016/j.jsb.2016.07.011.
15. Buchholz, T.O., Krull, A., Shahidi, R., Pigino, G., Jekely, G., and Jug, F. (2019). Content-aware image restoration for electron microscopy. Methods Cell Biol *152*, 277-289. 10.1016/bs.mcb.2019.05.001.
16. Liu, Y.T., Zhang, H., Wang, H., Tao, C.L., Bi, G.Q., and Zhou, Z.H. (2022). Isotropic reconstruction for electron tomography with deep learning. Nat Commun *13*, 6482. 10.1038/s41467-022-33957-8.
17. Chen, M., Bell, J.M., Shi, X., Sun, S.Y., Wang, Z., and Ludtke, S.J. (2019). A complete data processing workflow for cryo-ET and subtomogram averaging. Nat Methods *16*, 1161-1168. 10.1038/s41592-019-0591-8.
18. Winkler, H. (2007). 3D reconstruction and processing of volumetric data in cryo-electron tomography. J Struct Biol *157*, 126-137. 10.1016/j.jsb.2006.07.014.

19. Zhang, K. (2016). Gctf: Real-time CTF determination and correction. J Struct Biol *193*, 1-12. 10.1016/j.jsb.2015.11.003.
20. Morado, D.R., Hu, B., and Liu, J. (2016). Using Tomoauto: A Protocol for High-throughput Automated Cryo-electron Tomography. J Vis Exp, e53608. 10.3791/53608.
21. Heebner, J.E., Purnell, C., Hylton, R.K., Marsh, M., Grillo, M.A., and Swulius, M.T. (2022). Deep Learning-Based Segmentation of Cryo-Electron Tomograms. J Vis Exp. 10.3791/64435.
22. Schuck, P. (2000). Size-distribution analysis of macromolecules by sedimentation velocity ultracentrifugation and lamm equation modeling. Biophys J *78*, 1606-1619. 10.1016/S0006-3495(00)76713-0.
23. Philo, J.S. (2023). SEDNTERP: a calculation and database utility to aid interpretation of analytical ultracentrifugation and light scattering data. Eur Biophys J *52*, 233-266. 10.1007/s00249-023-01629-0.
24. Jumper, J., Evans, R., Pritzel, A., Green, T., Figurnov, M., Ronneberger, O., Tunyasuvunakool, K., Bates, R., Žídek, A., Potapenko, A., et al. (2021). Highly accurate protein structure prediction with AlphaFold. Nature *596*, 583-589. 10.1038/s41586-021-03819-2.
25. Abramson, J., Adler, J., Dunger, J., Evans, R., Green, T., Pritzel, A., Ronneberger, O., Willmore, L., Ballard, A.J., Bambrick, J., et al. (2024). Accurate structure prediction of biomolecular interactions with AlphaFold 3. Nature *630*, 493-500. 10.1038/s41586-024-07487-w.
26. Clementi, C., Nymeyer, H., and Onuchic, J.N. (2000). Topological and energetic factors: what determines the structural details of the transition state ensemble and "en-route" intermediates for protein folding? An investigation for small globular proteins. J Mol Biol *298*, 937-953. 10.1006/jmbi.2000.3693.
27. Latham, A.P., and Zhang, B. (2021). Consistent Force Field Captures Homologue-Resolved HP1 Phase Separation. J Chem Theory Comput *17*, 3134-3144. 10.1021/acs.jctc.0c01220.
28. Regy, R.M., Thompson, J., Kim, Y.C., and Mittal, J. (2021). Improved coarse-grained model for studying sequence dependent phase separation of disordered proteins. Protein Sci *30*, 1371-1379. 10.1002/pro.4094.
29. Eastman, P., Swails, J., Chodera, J.D., McGibbon, R.T., Zhao, Y., Beauchamp, K.A., Wang, L.P., Simmonett, A.C., Harrigan, M.P., Stern, C.D., et al. (2017). OpenMM 7: Rapid development of high performance algorithms for molecular dynamics. PLoS Comput Biol *13*, e1005659. 10.1371/journal.pcbi.1005659.
30. Liu, S., Wang, C., Latham, A.P., Ding, X., and Zhang, B. (2023). OpenABC enables flexible, simplified, and efficient GPU accelerated simulations of biomolecular condensates. PLoS Comput Biol *19*, e1011442. 10.1371/journal.pcbi.1011442.
31. Dignon, G.L., Zheng, W., Best, R.B., Kim, Y.C., and Mittal, J. (2018). Relation between single-molecule properties and phase behavior of intrinsically disordered proteins. Proc Natl Acad Sci U S A *115*, 9929-9934. 10.1073/pnas.1804177115.
32. Schrimpf, W., Barth, A., Hendrix, J., and Lamb, D.C. (2018). PAM: A Framework for Integrated Analysis of Imaging, Single-Molecule, and Ensemble Fluorescence Data. Biophys J *114*, 1518-1528. 10.1016/j.bpj.2018.02.035.

33. Hellenkamp, B., Schmid, S., Doroshenko, O., Opanasyuk, O., Kühnemuth, R., Rezaei Adariani, S., Ambrose, B., Aznauryan, M., Barth, A., Birkedal, V., et al. (2018). Precision and accuracy of single-molecule FRET measurements-a multi-laboratory benchmark study. Nat Methods *15*, 669-676. 10.1038/s41592-018-0085-0.
34. Kudryavtsev, V., Sikor, M., Kalinin, S., Mokranjac, D., Seidel, C.A., and Lamb, D.C. (2012). Combining MFD and PIE for accurate single-pair Förster resonance energy transfer measurements. Chemphyschem *13*, 1060-1078. 10.1002/cphc.201100822.
35. Tomov, T.E., Tsukanov, R., Masoud, R., Liber, M., Plavner, N., and Nir, E. (2012). Disentangling subpopulations in single-molecule FRET and ALEX experiments with photon distribution analysis. Biophys J *102*, 1163-1173. 10.1016/j.bpj.2011.11.4025.
36. Antonik, M., Felekyan, S., Gaiduk, A., and Seidel, C.A. (2006). Separating structural heterogeneities from stochastic variations in fluorescence resonance energy transfer distributions via photon distribution analysis. J Phys Chem B *110*, 6970-6978. 10.1021/jp057257+.
